# Quantitative analysis of population-scale family trees using millions of relatives

**DOI:** 10.1101/106427

**Authors:** Joanna Kaplanis, Assaf Gordon, Mary Wahl, Michael Gershovits, Barak Markus, Mona Sheikh, Melissa Gymrek, Gaurav Bhatia, Daniel G. MacArthur, Alkes L. Price, Yaniv Erlich

## Abstract

Family trees have vast applications in multiple fields from genetics to anthropology and economics. However, the collection of extended family trees is tedious and usually relies on resources with limited geographical scope and complex data usage restrictions. Here, we collected 86 million profiles from publicly-available online data from genealogy enthusiasts. After extensive cleaning and validation, we obtained population-scale family trees, including a single pedigree of 13 million individuals. We leveraged the data to partition the genetic architecture of longevity by inspecting millions of relative pairs and to provide insights to population genetics theories on the dispersion of families. We also report a simple digital procedure to overlay other datasets with our resource in order to empower studies with population-scale genealogical data.

**One Sentence Summary:** Using massive crowd-sourced genealogy data, we created a population-scale family tree resource for scientific studies.

---

Family trees are mathematical graph structures that capture two fundamental processes: mating and parenthood. As such, the edges of the trees represent potential transmission lines for a wide variety of genetic, cultural, socio-demographic, and economic factors in the human population. The foundation of quantitative genetics is built on dissecting the interplay of these factors by overlaying data on family trees and analyzing the correlation of various classes of relatives (*1*–*3*). In addition, family trees can serve as a multiplier for genetic information by using study designs that leverage genotype or phenotype data from relatives (*4*–*7*), analyzing parent-of-origin effects (*8*), refining heritability measures (*9*, *10*), or improving individual risk assessment (*11*–*13*). Beyond classical genetic applications, large-scale family trees have played an important role in a wide array of disciplines including human evolution (*14*, *15*), anthropology (*16*), and economics (*17*, *18*).

Despite the range of applications, constructing population-scale family trees has been a labor-intensive process. So far, the leading approach has mainly relied on local data record repositories such as churches or vital record offices (*19*–*21*). While playing an important role in previous studies, this approach has several limitations (*22*, *23*): first, it requires non-trivial resources to digitize the records and organize the data. Second, the resulting trees are usually limited in scope to the specific geographical area of the repository, which precludes comparisons between regions. Finally, vital record data is usually subject to various privacy protections and regulations. This reduces its accessibility and complicates its fusion with other sources such of information such as genomic or health data.

Here, we leveraged genealogy-driven social media data to construct population-scale family trees and create a layer of genealogy information for various studies. To this end, we focused on Geni.com, one of the largest crowd-sourcing websites in the genealogy domain. After careful cleaning of the tree data using various algorithms, we were able to obtain high quality datasets about family structure together with quantitative biographic information on individuals. We harnessed this massive information to dissect the genetic architecture of longevity and analyze familial dispersion processes in light of previous population genetics theories. The entire cleaned datasets are publicly available on FamiLinx.org for academic research. Beyond static use of the de-identified data, we describe a digital mechanism that enables researchers to dynamically fuse the family trees of our resource with data obtained from their own studies. This can enable the analysis of population-scale genomic or phenotypic datasets in the context of family trees.

## Constructing and validating population scale family trees

The Geni.com website allows genealogy enthusiasts to upload their family trees and create profiles for each individual. These profiles can include name, basic demographic information, and a photo (Fig. 1A). The website automatically scans new profiles to detect similarities to existing ones and when a match is detected, the website suggests the option to merge the profiles. By merging, the genealogists can create larger family trees beyond their individual knowledge and collaboratively co-manage profiles to improve their accuracy.

**Fig. 1.**
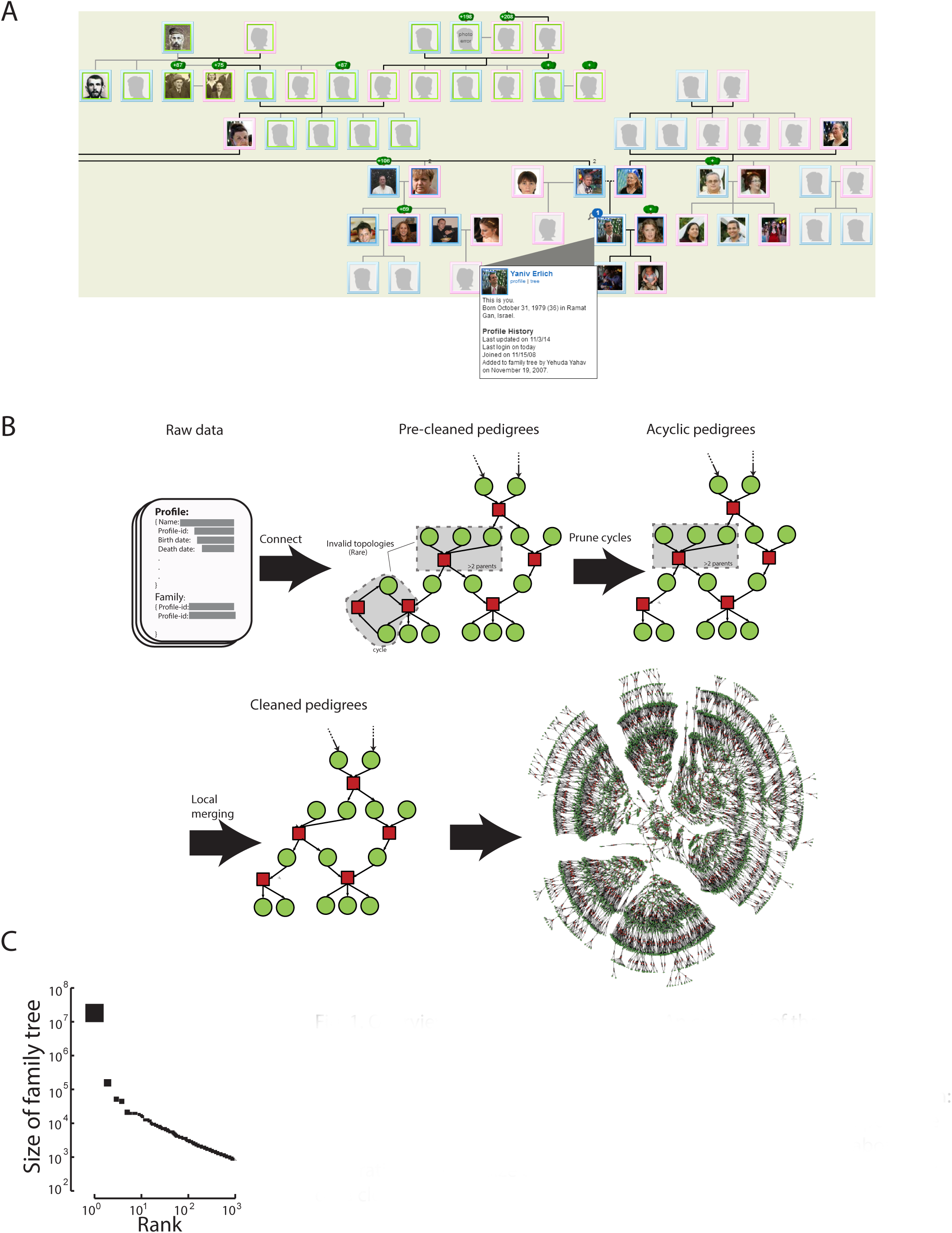
Overview of the collected data (A) An example of the genealogical and demographic information available on the website (B) The basic algorithmic steps to form valid pedigree structures from the input data available via the Geni API. See fig. S2 for a comprehensive overview. Green: profiles, red: marriages. The last step shows an example of a real pedigree from the website with ~6000 individuals. The family tree spans about 7 generations (C) The size distribution of the largest 1,000 family trees after data cleaning.

After obtaining IRB approval and permission from Geni and its parent company MyHeritage, we downloaded over 86 million publicly available profiles. This process required substantial data processing and organization, and took about six months to complete (*24*), but the underlying information is accessible to any Internet user and represents only public data that users have decided to share. The input data consists of millions of individual profiles, each of which describes a person and any putative connections to other individuals in the dataset, along with some auxiliary data about the creator of the profile. We found that more than 3 million genealogists took part in this large-scale crowdsourcing of family trees. Similar to other crowdsourcing projects (*25*), a small group of participants contributed most of the genealogy profiles (Fig. S1).

We organized the profiles into graph topologies that preserve the genealogical relationships between individuals (Fig. 1B) (*24*). Biology dictates that a family tree should form a directed acyclic graph (DAG) where each individual has an in-degree that is less than or equal to two. However, as expected from a large-scale collaborative work, 0.3% of the profiles resided in invalid biological topologies that included cycles (e.g. a person that is both the parent and child of another person) or an individual having more than two parents. We developed an automatic pipeline that used a series of graph-algorithmic steps and record-matching techniques to resolve local conflicts and prune invalid topologies (Fig. S2). To benchmark the performance of the pipeline, we compared the results with thousands of merging decisions by human genealogists and found high (>90%) concordance between the pipeline and human decisions regarding resolving conflicts (*24*). This process generated 5.3 million disjoint family trees.

The largest family tree in the processed data spanned 13 million individuals who were connected by shared ancestry and marriage (Fig. 1C). The tree included public figures such as Kevin Bacon and Sewall Wright. On average, the tree spanned 11 generations between each terminal descendant and their founders (Fig. S3). To the best of our knowledge, this is the largest family tree available for scientific analysis. The size of this pedigree is not surprising: familial genealogies coalesce at a logarithmic rate compared to the size of the population (*26*, *27*).

We sought to evaluate the structure of the tree by inspecting the genetic segregation of unilineal markers. We obtained mitochondria (mtDNA) and Y-STR haplotypes to compare multiple pairs of relatives in our graph (*24*). The mtDNA data spanned 211 lineages that included in total 1768 meiosis events (i.e. graph edges), whereas the Y-STR data was available for 27 lineages that spanned in total 324 meiosis events. Using a prior of no more than a single non-paternity event per lineage, the non-maternity rate was 0.3% per meiosis and the non-paternity rate was 1.9% per meiosis. Importantly, this rate of non-paternity closely matched previous rates in clinical studies (*28*, *29*). Taken together, these results indicate that the collaborative process of millions of genealogists can produce high quality family trees.

## Extracting high quality demographic data

Encouraged by the quality of the family trees, we extracted demographic information from the collected profiles (*24*). First, we analyzed the lifespan by focusing on profiles with full birth and death dates that show higher quality of data compared to profiles with only information about the year of birth (Fig. S4). Fig. 2A presents the overall distribution of age of death in the data with the exact birth and death dates. The data reflected well-known historical events and trends such as elevated death rates at military age the American Civil War, WWI, and WWII, and a reduction in child mortality during the 20^th^ century.

**Fig. 2.**
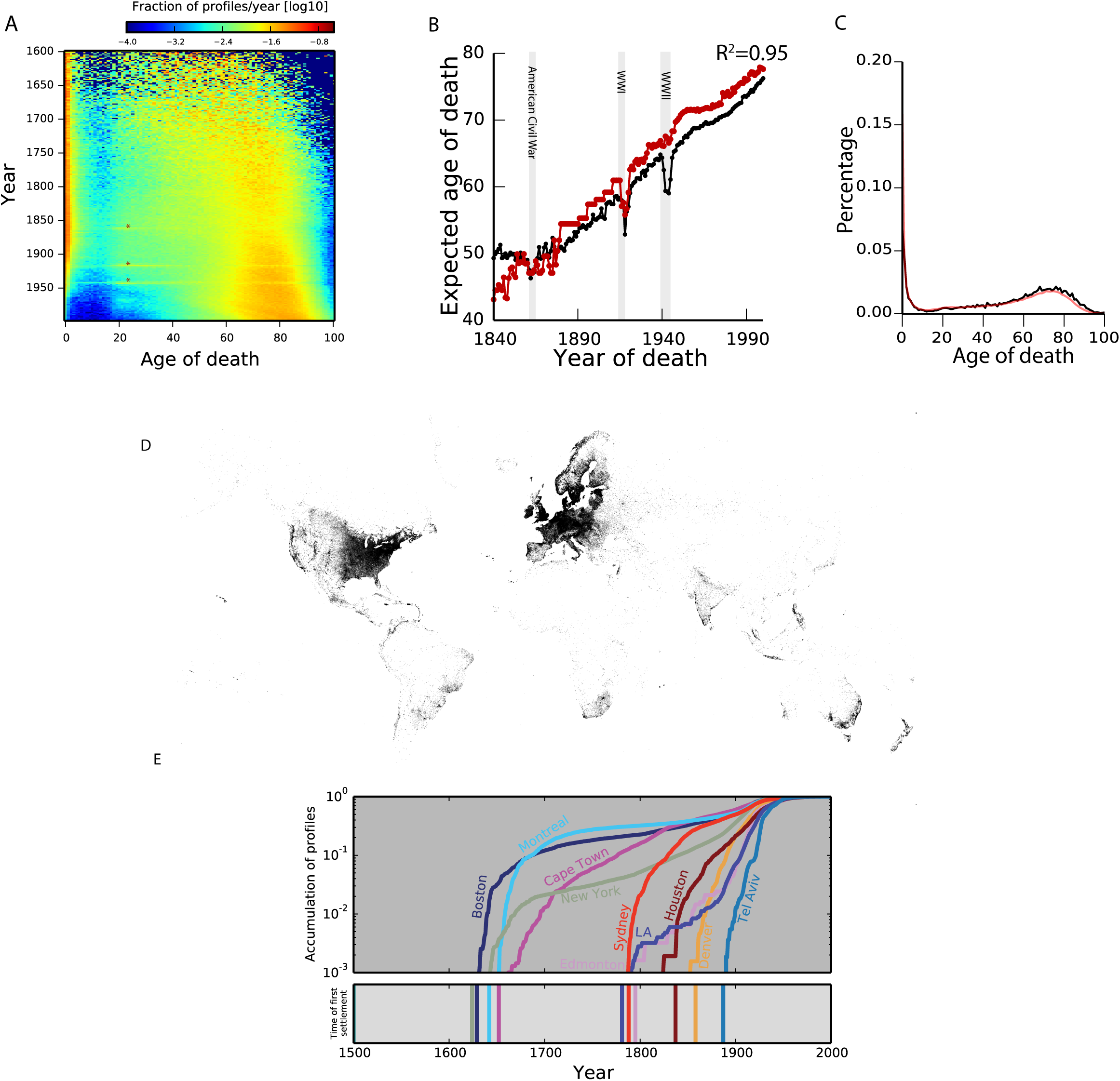
Analysis and validation of the demographic data (A) An overview of lifespan dynamics. The colors correspond to the fraction of profiles with a death reported at a certain age per year (B) The expected lifespan in our dataset (black) and the Oeppen & Vaupel study (red) as a function of year of death. The two major datasets are highly correlated (C) Comparing the lifespan distributions versus Geni (black) and HMD (red). Data is shown for 1880. Additional comparisons are in fig. S5A. (D) The geographic distribution of the annotated place of birth information. Every pixel corresponds to a profile in the dataset. The figure shows extensive sampling of the Western world (E) Validation of the geographical assignment by historical trends. Top: the cumulative distribution of profiles since 1500 for each city on a logarithmic scale as a function of time. Bottom: year of first settlement in the city. The color matches the city in the upper panel. Concordant with expectation, in all cases besides Houston and Denver, the accumulation of profiles starts immediately on or after the year of settlement.

The extracted lifespan information displayed excellent concordance with previous reports by traditional demographic approaches. We compared the average lifespan in our collection to a worldwide analysis based on historical data compiled from multiple countries around the world for each year from 1840-2000 (*30*). We found an R^2^=0.95 between the life expectancies in historical data and our dataset (Fig. 2B). As another layer of validation, we contrasted the lifespan distributions in our collection with the historical distributions reported by the Human Mortality Database (HMD) (*31*), which relies on vital records. The two datasets showed 98% concordance after inspecting the age of death distributions for 1820-1940 in intervals of 30 years (Fig. 2C; Fig. S5A). The only detectable systematic difference was an approximately 50% reduction in the mortality rate before the first birthday in the Geni data (Fig. S6B), presumably reflecting the inherent difficulty for genealogists of obtaining documentation of perinatal deaths that occurred several generations ago.

Next, we extracted the locations of life events using an automatic pipeline (*24*). Genealogists in the resource specified the birth and death locations as free text in various languages for tens of millions of profiles. We used geoparsing tools to annotate the free text into longitude/latitude coordinates. This process successfully reported the location of about 16 million profiles, typically at a fine-scale geographic resolution such as a town. The geographical distribution of the profiles showed a dense sampling of a wide range of locations in the Western World (Fig. 2D; fig. S6). About 30% of the profiles came from North America, 50% from Western Europe and Scandinavia, 5% from the British Isles, and the remaining profiles mapped to more than 100 other countries across the globe.

We assessed the overall accuracy of the geographic assignments by inspecting their concordance with historical events. To this end, we analyzed the accumulation of profiles in 10 major cities around the globe whose years of first settlement are known and occurred after 1600 (Fig. 2E). With an error-free geographic and time assignment, cities should have no profiles dated before the first known settlement. Consistent with this expectation, in nearly all of the examined cases, profiles appeared in the city only after the time of first settlement. In Houston and Denver only, about 0.2% of the profiles appeared before the existence of the city. **Movie S1** presents the place of birth of individuals in our data in 5 year intervals from 1400 to 1900 along with known migration events.

Taken together, our analyses show that accurate demographic information can be obtained from the cohort. Table S1 reports key demographic and genetic attributes for various familial relationships from parent-child via great-great-grandparents to fourth cousins. These include birth location distance, generation time difference, and the measured identity-by-descent (IBD) based on all genealogical ties.

## Characterizing the genetic architecture of longevity

We leveraged our data to characterize the genetic architecture of human longevity, a trait with profound public interest that exhibits complex genetics that are likely to involve a range of physiological and behavioral endophenotypes (*32*, *33*). Twin and nuclear family studies have estimated the narrow-sense heritability (h^2^) of longevity to be around 15%-30%, with 25% as the typically cited value in the literature (Table S2) (*34*–*39*). To date, genome-wide association studies have had very limited success in identifying genetic variants that are associated with longevity (*40*–*42*). This relatively large proportion of missing heritability can stem from the following non-mutually exclusive explanations: (A) longevity has substantial non-additive components (such as dominance or epistasis) that create upward bias in estimates of heritability (*43*), (B) previous estimators of heritability are upward biased due to environmental effects that were not properly accounted for (*10*), (C) the trait is highly polygenic and requires larger cohorts to identify the underlying variants (*44*). By harnessing our resource, we sought to build a model for the sources of genetic variance in longevity that jointly evaluates additivity, dominance, epistasis, shared household effects, spatiotemporal trends, and random noise.

To prepare the data for genetic analysis, we first adjusted spatiotemporal effects (*24*). We evaluated a series of nested models with a range of non-genetic factors that have been associated with longevity, namely sex, year of birth, country of birth, the exact longitude and latitude of birth location, and the average temperature. The training set for these models consisted of approximately 3 million individuals from our resource who had exact data regarding date of birth, death, and location, and lived at least to the age of thirty, as our motivation was to study lifespan during adulthood. For validation, the models reported their goodness of fit (R^2^), mean squared error, Bayesian Information Criterion (BIC), and the statistical significance of each factor using an independent set of 300,000 individuals. The best model included sex, birth year, and the exact longitude and latitude (Fig. 3A); simpler models had higher MSE, lower R^2^, and worse BIC levels, while more complex models that included the country of birth or the temperature did not improve the goodness of fit, and the additional factors did not reach statistical significance in most cases. The best model explained about 7% of the longevity in the validation set. We defined longevity as the difference of the age of death from the expected lifespan based on the best model.

**Fig. 3.**
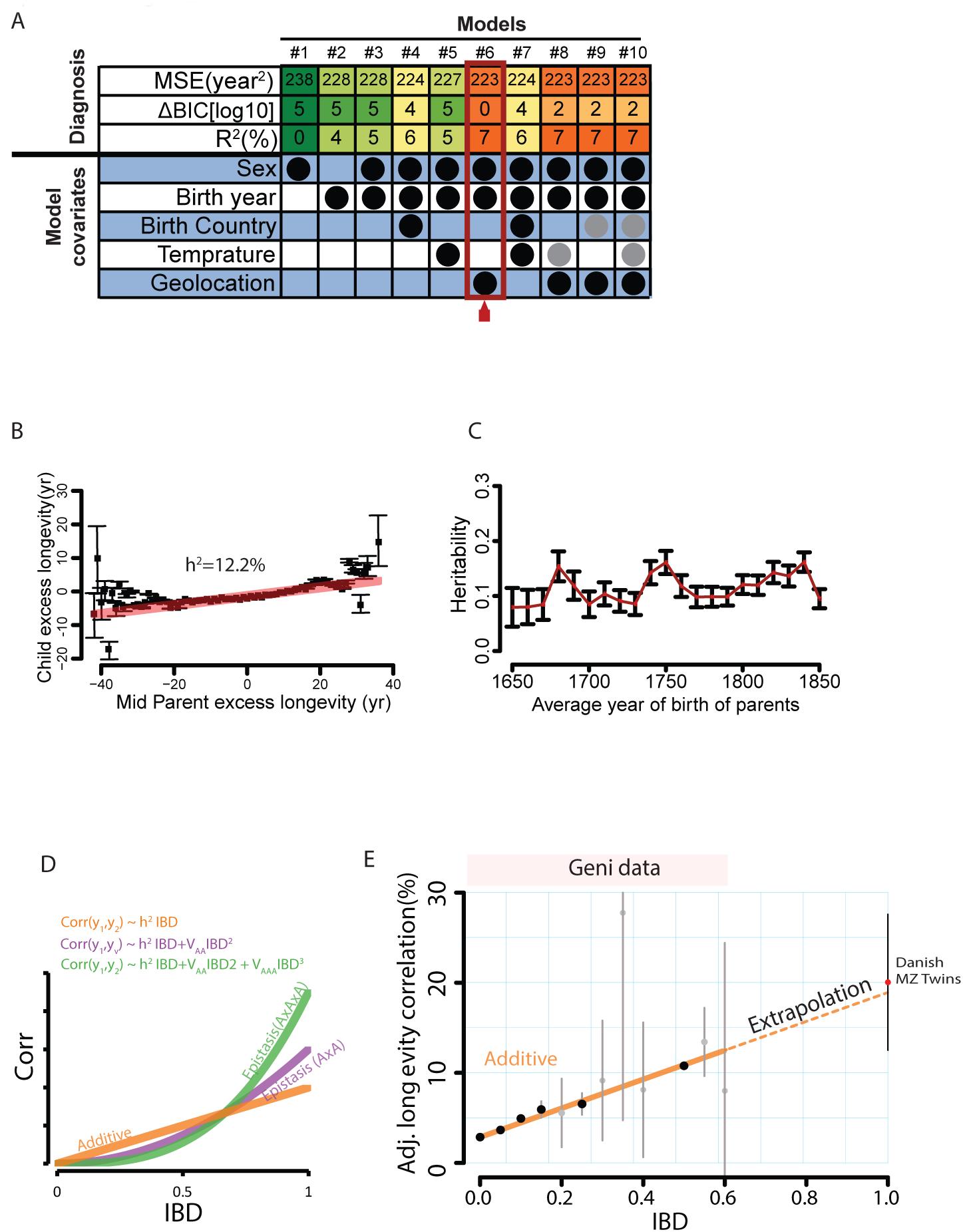
The genetic architecture of longevity. (A) Adjustment of longevity using various environmental models. MSE: mean squared error per individual; ΔBIC is the difference of the Bayesian Information Criterion from the best model after log10(x + 1) transformation. Circles represent covariates in each model. Gray/Black: statistically in/significant (p < 0.01) covariate. Orange to green: most desired to least desired diagnostic outcome. The best model is #6, which adjusts longevity based on sex, year of birth, and geolocation. This model had the lowest MSE, smallest BIC, and best R^2^ (B) The regression (red) of child longevity on its mid-parent longevity (defined as difference of age of death from the expected life span). Black: the averaged longevity of children binned by the mid-parent value. Gray: estimated 95% confidence intervals (C) The estimated narrow-sense heritability (black squares) with 95% confidence intervals (black bars) obtained by the mid parent design stratified by the average decade of birth of the parents (D) The Kempthorne model describes the expected correlation between relatives for a quantitative trait as a function of their genetic resemblance. For simplicity, the figure shows the correlation of a trait as a function of IBD under strict additive (h^2^, orange), squared- (V_AA_, purple), and cubic- (V_AAA_, green) epistasis architectures (E) The average longevity correlation as a function of IBD (black circles) grouped in 5% increments (gray: 95% CI) after adjusting dominancy. The epistatic terms converged to zero and therefore are identical to the presented additive model. Dotted line: the extrapolation of the models towards MZ twins from the Danish Twin Registry (red circle).

We sought to further validate the quality of our data by estimating the narrow-sense heritability of longevity (*h*^2^) according to the mid-parent design (*45*, *46*). We analyzed nearly 130,000 parent-child trios with the highest quality data (exact date of birth, death, and town resolution for place of birth) (*24*). This process yielded *h*^2^_mid-parent_ = 12.2% (s.e.=0.4%) (Fig. 3B), which falls within the range of previous heritability estimates, but on the lower end (Table S2). The data did not show any significant linear correlation (p>0.1) between heritability and the decade of birth of the parents (Fig. 3C), matching previous findings (*35*, *37*). We also did not find any significant difference (p>0.4) between the heritability of longevity from mother-offspring pairs (*h*^2^_mother_ = 12.8%, s.e.=0.4%, n=220,000) versus father-offspring pairs (*h*^2^_father_ = 13.2%, se=0.4%, n=271,000). However, we did observe a significant difference (p<10^−11^) between the heritability of concordant-sex parent-offspring pairs (e.g. mother-daughter) versus discordant-sex pairs (e.g. mother-son), with *h*^2^_concordant/parent-child_ = 15.0% (se=0.4%, n=254,000) and *h*^2^-discordant/parent-child = 10.7% (s.e.=0.4%, n=236,000). This difference most likely arose due to differences in the distribution of longevity between sexes; women show higher mortality than men around child bearing ages but lower mortality after these ages (Fig. S7). Our model only adjusts for the average life expectancy of each sex and we posit that the reduced *h*^2^_discordant/parent-child_ stems from the differences between the distributions.

We partitioned the source of genetic variance of longevity using over three million pairs of relatives from full sibling to 4^th^ cousins (Table S1) (*24*). These pairs were obtained from a core set of nearly half a million individuals that passed all inclusion criteria and have precise demographic data regarding their place of birth and longevity information. To mitigate correlations due to non-genetic factors, these three million pairs do not include relatives that are likely to have died due to environmental catastrophes or in major wars (Fig. S8) and consisted of only sex-concordant pairs to mitigate residual sex-differences that were not accounted for by our longevity adjustments. For each pair of relatives, we calculated the Jacquard’s nine condensed identity coefficients of the pairs (*47*) in order to include in the IBD calculation the effects of historical inbreeding (Fig. S9-**10**). After that, we evaluated the dominance variance by inspecting the difference in the longevity correlation of full siblings versus father-son pairs using over 300,000 pairs of relatives. Then, we adjusted the dominance variance and used the Kempthorne model (*48*) to decompose the contribution of additive genetic variance, quadratic epistasis variance, and cubic epistasis variance (Fig. 3D).

The analysis of longevity in these 3 million of pairs of relatives found a robust additive genetic component, small impact of dominance, minimal household effects, and no evidence of epistasis (Fig. 3E; Table S3) (*24*). Additivity was highly significant (p_additive_<10^−318^) with an estimated *h*^2^_concordant/relatives_ = 16.1% (s.e.=0.4%), similar to the heritability estimated from sex-concordant parent-child pairs *h*^2^-_concordant/parent-child_ = 15.0% (s.e.=0.4%). The estimated dominance was 4.0% using the father-son and full sib pairs and the shared household effect explained nearly 1% of the overall correlation. Despite the substantial amount of data with 3 million pairs of relatives, the quadratic epistasis factor and the cubic epistatic terms converged to zero (quadratic s.e.= 0.1%, cubic s.e.=0.1%). Bayesian Information Criterion (BIC) also strongly disfavored the quadratic epistatic factor (*ΔBIC* = 15) and the cubic factor (*ΔBIC* = 30). We also tested a cross validation procedure that repeatedly removed classes of genealogical relationships (e.g. second cousins), fit the data with and without epistatic terms, and measured the MSE between the model prediction and the removed class. This procedure did not decrease the MSE when epistatic terms were present, again arguing against their contribution to longevity variance in the population.

To further validate these results, we tested the ability of our model to predict the longevity correlation of an orthogonal dataset of monozygotic (MZ) twin pairs collected by the Danish Twin Registry (Fig. 3E) (*49*). This dataset of 810 MZ twins was collected by traditional epidemiological means and is highly homogenous in terms of time (birth between 1870-1900), spatial dispersion, and reflects one of the most socioeconomic equal countries in the world. Strikingly, we found that our inferred model for longevity accurately predicted the observed correlation of the twin cohort with 1% difference, which is well within the sampling error for the mean twin correlation (s.e. = 3.2%). We also repeated the analysis by adding the twin data to the Geni dataset and analyzing the entire collection for quadratic epistasis. Again, the quadratic epistasis converged to zero (95% CI: 0-2.7%). As interaction terms have maximal contribution to the resemblance of MZ twins, the consistency of the additive model confirms their marginal role. Furthermore, it also limits the role of various types of complex interactions that were not included in our model such as additive-dominant interactions or higher order epistatic interactions. Importantly, these results also demonstrate the power of our social media-derived dataset to accurately predict the findings of a study collected by conventional means, providing an extrinsic validation to the overall approach. Finally, we repeated the analysis with more complex adjustments including further correcting for household effects (fig. S11-**12A,B**), addressing potential errors in family trees (fig. S12C), or removing possible confounders due to inbreeding (Fig. S12D) (*24*). Yet, the results remained highly consistent (Fig. S12e). Additivity explained 15.8%-16.9% of the longevity estimates, dominance explained 2%-4%, the household environment contributed about 1%, and we did not find any sign of epistatic interactions.

## Assessment of population genetics theories of familial dispersion

Familial dispersion is a major driving force of various genetic, economical, and demographic processes (*50*). Yet, high-grade datasets of successive generations of human migration are scarce. Previous work has primarily relied on vital records from geographically limited areas (*51*, *52*) or used indirect inference from genetic datasets [for review: (*53*)] that mainly illuminate far historical events. To demonstrate the power of our data, we harnessed our resource to evaluate previous hypotheses in the literature regarding patterns of human migration.

First, we analyzed sex-specific migration patterns (*24*). Previous studies using genetic data reported conflicting results regarding sex bias in human migration (*54*). Earlier results reported decreased mitochondria differentiation compared to Y-chromosome, suggesting higher migration events for females (*53*). However, this pattern was not found in later studies of larger geographic sampling and more advanced genotyping technology (*56*). To better understand these results, we inspected the birth locations of parents versus their offspring and stratified the trend by the sex of the parent and the year of birth (Fig. 4A [colored lines]).

**Fig. 4.**
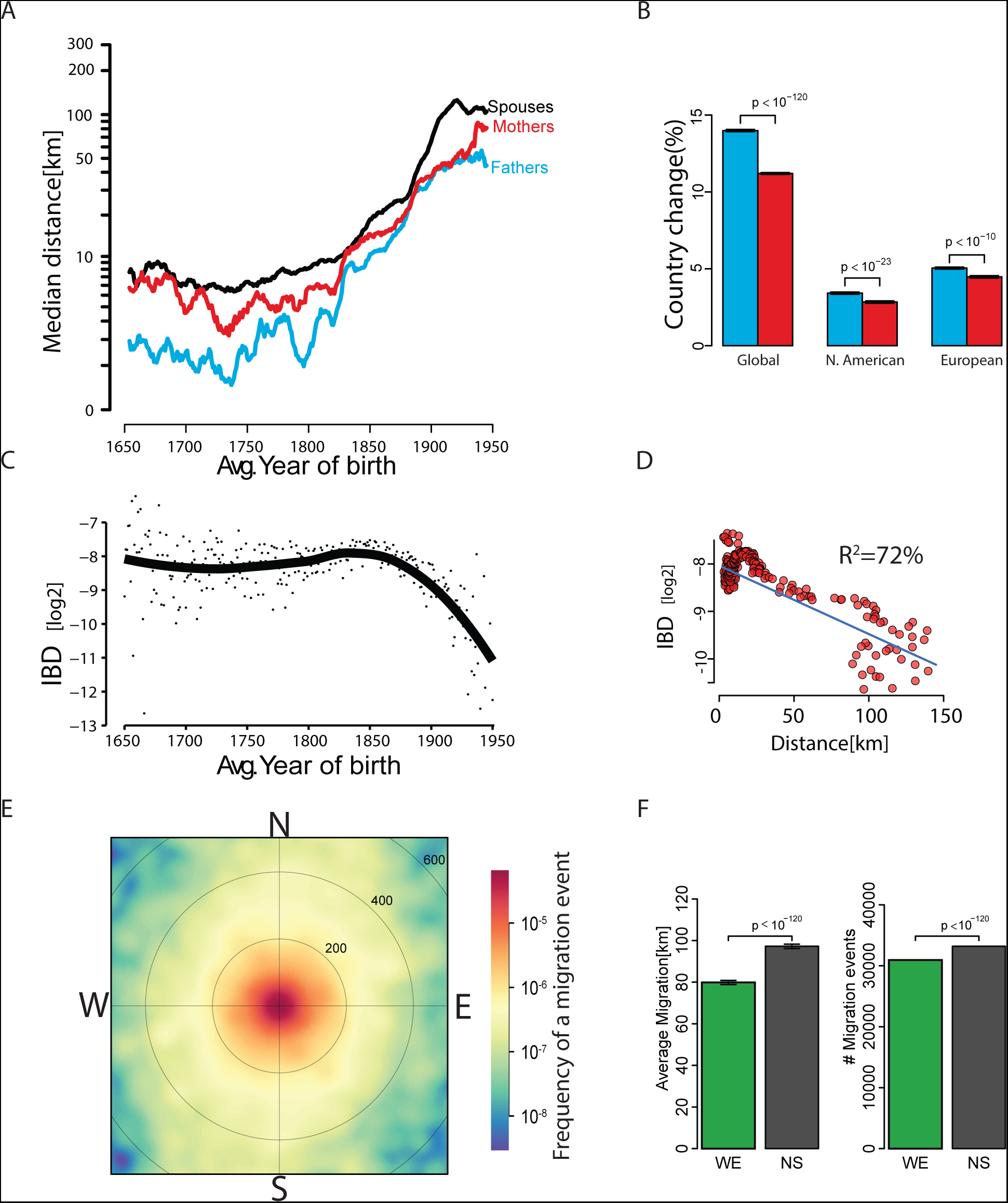
Analysis of familial dispersion. (A) The median distance [log_10_ x+1] of father-offspring places of birth (cyan), mother-offspring (red), and marital radius (black) as a function of time (average year of birth) (B) The rate of change in the country of birth in offspring vs. father (cyan) or mother (red) stratified by major geographic areas (C) The average IBD [log_2_] between couples as a function of average year of birth. Individual dots represent the measured average per year. Black line denotes the smooth trend using locally weighted regression (D) The IBD of couples as a function of marital radius. Blue line denotes best linear regression line (in log-log space) (E) Heat map of 60,000 migration events in Europe measured as the distance and bearing between parent and offspring. Circles denote the distance in Km (F) Analysis of migration azimuth strongly shows that NS migration events are more prevalent than WE events.

Our data indicates that females migrate more than males but over shorter distances in Western societies. The *median* mother-child distances were significantly larger (Wilcox, one-tailed, p<10^−90^) by a factor of 1.6x than father-child distances. This trend appeared throughout all 300 years of our analysis window including in the most recent birth cohort and was observed both in North American duos (Wilcox, one-tailed, p<10^−23^) and European duos (Wilcox, one-tailed, p<10^−87^). On the other hand, we found that the *average* mother-child distances (Fig. S13) were significantly shorter than the father-child distances (t-test, p<10^−90^), suggesting that long-range migration events are biased towards males. Consistent with this pattern, fathers displayed a significantly (p<10^−83^) higher frequency than mothers to be born in a different country than their offspring (Fig. 4B). Again, this pattern was evident when restricting the data to North American or European duos. Taken together, at least in the Western societies in our database, males and females show different migration distributions in which patrilocality occurs only in relatively local migration events and largescale events that usually involve a change of country are more common in males than females.

Next, we inspected the marital radius (the distance between mates’ places of birth) and its effect on the genetic relatedness of couples (*24*). The classical isolation by distance theory of Malécot predicts that changes to the marital radius should exponentially decrease the genetic relatedness of individuals (*57*). But the magnitude of these forces is also a function of technological and cultural factors such as taboos against cousin marriages (*58*). To understand the interplay between these factors, we first inspected temporal changes in the birth locations of couples in our cohort. Pre-industrial revolution (<1750), most marriages occurred between people born only 10km from each other (Fig. 4A [black line]). Similar patterns were found when analysing only European-born individuals (fig. S14) or North American-born individuals (Fig. S15). After the beginning of the second industrial revolution (1870), the marital radius rapidly increased and reached ~100km for the birth cohort of 1950.

We then inspected the genetic relatedness (IBD) of couples as measured by tracing their genealogical ties (Fig. 4C) (*24*). In 1650-1850, the expected IBD of couples was relatively stable and was in the order of ~4^th^ cousins, whereas post-1850, the IBD exhibited a rapid decrease. Overall, the median marital radius of each year showed a strong correlation (R^2^=72%) with the expected IBD between couples. Every 70km increase in the marital radius corresponded approximately with a decrease in the genetic relatedness of couples by one meiosis event (Fig. 4D), which matches previous isolation by distance forces in continental regions (*59*). However, this trend was not consistent over time and exhibits three phases. For the pre-1800 birth cohorts, the correlation between marital distance and IBD was insignificant (p>0.2) and weak (R^2^=0.7%) (Fig. S16A). Couples born around 1800-1850 showed a two-fold increase in their marital distance from 8km in 1800 to 19km in 1850. Marriages are usually about 20-25 years after birth and around this time (1820-1875) rapid transportation changes took place, such as the advent of railroad travel in most of Europe and the United States. However, the increase in marital distance was significantly (p<10^−13^) coupled with an *increase* in genetic relatedness, contrary to the isolation by distance theory (f**ig. S16B**). Only for the cohorts born after 1850, the data showed a good match (R^2^=80%) with the theoretical model of isolation by distance (Fig. S16C). Taken together, the data shows a 50-year lag between the advent of increased familial dispersion and the decline of genetic relatedness between couples. During this time, individuals continued to marry relatives despite the increased distance. Based on these results, we hypothesize that the marked transportation changes in the 19^th^ century were not the primary cause for decreased consanguinity. Rather, our results suggest that shifting cultural factors, such as change in attitude towards consanguinity, played a more important role in the reduced genetic relatedness of couples in Western societies since the 19^th^ century.

Finally, we wondered whether familial dispersion has a dominant direction. Diamond (*60*) has postulated that certain technologies such as agricultural strategies diffuse faster along the latitudes (east-west, WE) due to climate similarity, promoting human migration along the same axis. Indeed, previous genetic studies have found that Europeans show greater genetic differentiation in the NNW-SSW axis, suggesting increased migration along the perpendicular axis during the Paleolithic-Neolithic era (*61*, *62*). Consistent with these observations, a contemporary analysis showed an increase in linguistic diversity in countries that are more WE oriented than NS oriented (*63*). To measure the orientation of dispersion, we analyzed the birth locations of parent-offspring duos in our data (*24*). In order to remove the effects of the mass migration to the New World and the US gold rush, we only focused on European-born duos and required a minimum 10km separation between birth places to obtain a clear orientation vector (Fig. 4E).

Our data argue against WE preference in Europeans (Fig. 4F). The average migration along the NS axis was significantly longer than the WE axis (NS: 97km, WE: 80km, p<10^−120^). To gain more evidence, we repeated this analysis by counting the number of migration events with WE orientation versus NS orientation rather than the distance. Again, the rate of NS migration events was significantly larger than the WE events (p<10^−120^). Since climate is highly similar between adjacent locations, we were concerned that a 10km separation threshold is too small to observe orientation differences. We repeated the analysis with a 100km threshold. Also in this case, migrations along the NS axis were significantly higher than the WE axis, with a higher average migration distance (NS: 208km, WE: 159km) and the rate of preferred migration along the axis (NS: 58%, WE: 42%) (Fig. S17A). We also restricted the analysis to a minimal separation threshold of 1km for events to since the second industrial revolution (post-1850) (Fig. S17B) or a (Fig. S17C). In all cases, the analysis did not find any preference for the WE orientation. Finally, we repeated the entire analysis while considering the median migration distance instead of the mean. Again, we observe a preference towards NS migration rather WE. All of these trends show that Diamond’s postulation about WE preference is not relevant for Europe after the Middle Ages.

## Discussion

In this work, we leveraged genealogy-driven media to build a dataset of human pedigrees of unprecedented size that covers virtually every country in the Western world. Our multiple validation procedures indicate that it is possible by careful cleaning to obtain a highly reliable demographic dataset that has similar quality to traditionally collected studies, but with much larger scale and at lower cost.

Using these data, we dissected the genetic architecture of longevity. The consistency across conditions shows the robustness of the results and highlights that the genetic variance of longevity is mainly mediated via additive and dominant effects, with no detectable signal of gene-gene interactions (epistasis). Our results indicate that the limited ability of GWAS studies so far to associate variants with longevity cannot be attributed to statistical epistasis. We note that additive genetic architecture for human longevity does not contradict the existence of molecular interactions between genes contributing to this trait (*64*–*66*). Our analysis reflects the net effect of epistasis, integrating the frequency and impact of genetic variation of these molecular interactions. Depending on the frequency spectrum and the epistasis type, pervasive molecular interactions may have little to no epistatic contributions to phenotypic variance (*67*). Importantly, we also found that the additive component of heritability (*h*^2^_concordant/relatives_ = ~15%) measured in our dataset with millions of people is substantially lower than the value generally cited in the literature (25%). These results indicate that previous studies are likely to have overestimated the heritability of longevity. As such, we should reduce our expectations about our ability to predict longevity from genomic data and to identify causal genetic variants.

We also tested a series of hypotheses in population genetics regarding familial dispersion in humans. The data exhibit marked differences in the migration profiles of females and males and favored the hypothesis that, at least in Western societies, patrilocality operates only on a local scale. In addition, we used the data to investigate the isolation by distance theory in humans. This analysis suggests that the advent of the transportation revolution had a massive impact on familial dispersion but was not the primary cause of reduced consanguinity in recent generations. Rather, the data showed a period of nearly two generations of increased relatedness between couples despite elevated familial dispersion. Finally, we investigated Diamond’s hypothesis of east-west migration in Europe. Applying multiple analytical techniques, the data consistently demonstrated the opposite: most European migration in the last 300 years was actually north-south.

We envision that the data from our resource can address further questions in disciplines that are interested in quantitative aspects of human families, including genetics, anthropology, public health, and economics. Researchers can use the raw data in two ways. First, the entire tree and demographic data are available in a de-identified format. This enables static analysis of the datasets presented in this study. Second, we also offer a dynamic method that enables fusing other datasets with our data. Using the native Geni API, we created a simple web button that allows participants to identify themselves using their own account credentials, similar to the “login with Facebook” button (Fig. S18). Researchers can use this digital mechanism to obtain consent from a participant and collect the profile number of the participant. This will enable the connection between a real person and the genealogy data in our resource. The supplementary material (*24*) contains links to an example page and a GitHub repository with the code for the button. We currently use this one-click mechanism to overlay genomes with our family tree on DNA.Land (*68*). After consent, participants of DNA.Land simply enter their Geni.com username and password, which are sent directly to Geni.com, who authenticates them, and reveals the profile number of the participant to DNA.Land. As our data is totally open, other projects can use a similar strategy to add large pedigrees to their existing data collection efforts.

More generally, our work demonstrates the potential of using the growing world of social media and crowd-sourced genealogy to address key questions in human genetics. Currently, genealogy websites are limited to rather basic information and do not contain genome-wide genetic data. However, with ever-growing digitization of daily life and the advent of direct-to-consumer genetics (*69*), ethically aware approaches for harnessing these resources can be a valuable path to reach the dramatic scale of information needed for large-scale scientific studies.

## Acknowledgments

We thank MyHeritage.com and Geni.com for their assistance. This study was supported by a generous gift from Andria and Paul Heafy (Y.E), the Broad Institute’s SPARC: Catalytic Funding for Novel Collaborative Projects award (Y.E. and D.M.), the Burroughs Wellcome Fund Career Awards at the Scientific Interface (Y.E.), and by NIH grants R01 GM105857 and R03 HG006731 (A.L.P.). Some of the data used for this research were provided by the Danish Twin Registry, University of Southern Denmark. The findings, opinions and recommendations expressed therein are those of the authors and are not necessarily those of the DTR. The authors thank D. Zielinski and J. Novembre for valuable comments. Y.E wants to thank the constant support of the Erlich lab members in pursuing this project.

### Conflict of interest statement

Y.E is a paid consultant of MyHeritage.

## Supplemental Methods

### Data acquisition and cleaning

#### Initial round of data gathering

We built a custom Perl script based on Geni.com `RESTful APIv0` to systematically download public data from the website. This process took months due to the rate limitation of Geni.com on third party applications. The returned dataset was millions of `JSON` files that represented individual profiles and their immediate families (see an example of the public profile of Sewall Wright). We parsed the familial connections in the returned `JSON` files and represented the data as a graph using the `C++ Boost Graph Library` and custom `Python` scripts.

The topology of family trees can be represented as a bipartite directed graph *G* = (*P, U, E*), where the *P* nodes represent individuals (profiles on Geni.com), *U* nodes represent mating between two individuals, and *E* represents the set of edges between the nodes. Let *|X|* denotes the size of set *X*. *G*_*raw*_, initially the raw graph prior to any processing, had *|P|* = 43.72 million individuals, *|U|* = 13.3 union events, and 4.5 million connected components, each of which denotes a family tree. The largest connected component was a family tree of 15.4 million people.

#### Removing cycles

Biology dictates that *G* is a Direct Acyclic Graph (DAG) because an offspring cannot be the ancestor of any of his parents. Consanguinity does not create cycles, rather it induces multiple paths between an ancestor and an offspring. However, *G*_*raw*_ had cycles presumably due to errors and the fact that the data was collected over three months while active merging events have been carried by the Geni genealogists. To identify cycles in *G*_*raw*_, we used Tarjan’s strongly connected components algorithm (strong connected component is a subgraph where every node is reachable by any other node and contains one or more cycles) [13]. We found a total of 1018 strong components, the largest component included 3814 vertices (profiles and unions) and the average component had 15 vertices. The most common cycle was an offspring that is the parent of one of his parents. We removed all 15, 256 nodes that were part of the strong components and the edges that connected these nodes to the rest of the graph.

Cycle removal had a minimal effect on the overall topology of the graph. The number of individuals in the largest connected component was reduced by 0.3% to 15.3 million and the number of connected components was increased by the same percentage.

#### Cleaning up multi-parents

Since each reproductive event is generated by exactly two individuals and every individual is the product of a single reproductive event, the maximal indegree for any node in *U* cannot be larger than two and the maximal indegree for any node in *P* cannot be bigger than one. Again, we found nodes that violate this assertion and had more than two parents or were assigned to more than one union event. Our initial strategy was to eliminate these events when possible by locally merging profiles that were a mere duplication of each other (Figure S2).

First, we employed a merging up procedure. We created an algorithm that scanned for union nodes with indegree more than 2 or profile nodes with indegree more than 1. When such a node was identified, the algorithm retrieved its parents and evaluated whether multiple parental nodes were referring to the same person. For that, the algorithm employed a similarity test for each pair of parents. The test consisted of checking whether the reported sex of the two parental profiles were concordant and whether their first names matched under the phenotypic Soundex system which can accommodate spelling variants. If the two profile nodes passed the similarity test, the algorithm merged them by transferring the edges of one node to the other node and excluding the former node from the dataset. Next, the algorithm moved one level up and merged the upstream union nodes by transferring the edges from one node to the other. The algorithm kept moving upwards towards the ancestral nodes like a zipper, performing similarity tests to putative profile nodes and merging them and their upstream union nodes. In order to increase the reliability of the merges, the algorithm committed to the merges only if it did not fail in any similarity test along 5 successive generations from the initial merge. Otherwise, the algorithm undid all operations originated in this round, restoring this section of the graph to its original topology.

Next, we employed a merging down procedure. The algorithm scanned again the nodes that were merged in the previous step but this time in went downwards in order to merge duplicate profiles of descendants. Since the algorithm already decided that there are two duplicated family trees, it just employed the similarity test for each generation until no more profiles were available.

Before the merging procedure, there were 354, 502 nodes with more than two parents. The merging up procedure merged 321, 985 nodes (including ancestral nodes to the one that triggered the procedure) and the merging down procedure merged 132, 623 nodes. In total, the algorithm resolved 180, 507 out of the 354, 502 nodes that had more than two parents.

The last step of the algorithm was to discard nodes with more than two parents that could not be resolved with the local merging procedure. Similar to the cycle clean up, we excluded those nodes from the dataset and pruned the edges connecting them to the rest of the graph.

The clean-up step had an effect on the overall topology of the graph. The number of individuals in the largest connected component was reduced by 15% to 13 million and the number of connected components was increased by 1.17%.

#### Estimating clean up accuracy

To evaluate the accuracy of the merging procedure, we compared the decisions of the merging algorithm to the decisions of the Geni.com genealogists. We randomly selected 1000 profile pairs that were merged during the merging up step and downloaded them again from the Geni.com website after some time from the initial collection of the data. We found that in 966 cases, the genealogists merged these profiles with each other or simply deleted one of them. In 18 cases, one of the profiles was merged to another profile but not to the one that was predicted by the merging algorithm, and in 16 cases no action was taken. The last two cases represent either potential failures of the merging algorithm or just incomplete work of the genealogists. Therefore, we estimate that at least 96.6% of the up-merges were correct based on agreement with the genealogists decisions.

We employed the same procedure for the down-merges. Here, 84% of the merges by our algorithm were concordant with the the decisions made by the genealogists. The size of the ‘no action’ cases was much larger (65 cases). We presume that the lower concordance in the down merges is in part attributed to a lack of a clear trigger for merging for the genealogists. It is easier for the genealogists to see more than two parents in their shared trees, which will initiate a merging up event. However, the genealogists do not always finish the process and eliminate duplicated nodes from the newly merged trees.

To get an additional independent estimator of the merging concordance, we also examined the profile photos that were associated with pairs of merged nodes. The hypothesis of this procedure was that profiles that are true duplicates of each other will have profile photos of the same person. We asked two human raters to independently inspect pairs of photos of profiles that were merged and to report whether the same person appears in both photos. To increase the reliability of the analysis, we included only cases where the reports of the two raters was identical. In 161 out of 171 merge cases with pairs of facial photos, the two raters concluded that the same individual appeared in both photos. With this result, the correct merging rate is estimated to be around 94%.

#### Second round of data collection

The initial round of data collection was done in 2011. In 2015, we scanned the website again to update our dataset. This round increases the size of our collection to 86 million unique profiles. We merged this new profiles with the old using the profile-id numbers of Geni. When the genealogists merge two profiles together, the Geni data flags one of the profiles a obsolete and indicated the newly consolidated profile (”merge-to”).

We followed all “merge-to” events and consolidated the information of the profiles in our clean tree. Using this procedure, we took the most up to date data and copied to the clean trees. To estimate the number of genealogists that contributed the data, we used the “created-by” field. All data was stored in a PostgreSQL database and is available on the FamiLinx website.

### Estimating the accuracy of the tree using genetic markers

We sought to evaluate the quality of the pedigree data in Geni.com using genetic markers. To this end, we downloaded publicly available records that include (a) Y chromosome STR markers or mitochondrial D-loop markers. Again, we only obtained data that is publicly available to any user with Internet connection and were voluntarily shared by individuals.

If two individuals are truly related, the distribution of the number of mismatches in their genetic markers can be approximated as a Poisson process with a parameter *λ* = *µ* · *g* · *n*, where *µ* is the mutation rate per locus per meiosis, *g* is the total number of meiosis events between the records, and *n* is the number of loci that were considered. Following previous studies [14], we set *µ_Y−STR_ ≈* 1*/*500 mutations per meiosis per marker and *µ_mito_ ≈* 3 × 10^*−*5^ mutations per meiosis per nucleotide.

The distribution of the number of mismatches between individuals that are not related was calculated by taking random pairs of records in our dataset and empirically measuring the number of mismatches.

We determined for each pairs of records the number of meiosis events between them and the observed number of mismatches and compared the two hypotheses whether the individuals are likely to be related or not. For that, we approximated the two distributions of the number of mismatches above as Gaussians. Next, we calculated a likelihood ratio test with an equal prior to discriminate between the hypotheses that the pair of individuals is related versus unrelated.For the mitochondrial data, we were able to test 209 pairs of allegedly related individuals through their maternal lines, which spanned 1,768 meiosis events. Only five pairs of individuals were unrelated according to the likelihood ratio test. As the prior expectation for non-maternity is low, we assumed a single non-maternity event per erroneous maternal line. With that, the maximum likelihood rate of non-maternity per is 5/1768 = 0.3% per meiosis event. For Y-STR data, we were able to test 28 pairs of allegedly related individuals, which spanned 324 meiosis events. Six pairs of individuals were unrelated according to the same procedure. Assuming a single non-paternity event per erroneous line, the maximum likelihood rate of non-paternity rate was 6/324 = 1.9% per meiosis event.

The rate of non-paternity matches previous estimates of Europeans. A meta-analysis based on 67 studies estimated the non-paternity rate to be 2% in Europeans [3]. More recently, a surname study in the UK with over 1,500 samples found a quite similar non-paternity rate [8]. In summary, the concordance of the non-paternity rates provide further support for the quality of the social media data.

### Extracting longevity information

#### Obtaining age of death

The year of birth and death for each profile was extracted by using the `birth_date` and `death_date` fields. Full date was defined as (i) entries that have non-empty strings in their corresponding month, day, and year fields AND (ii) the “circa” flag in the Geni JSON is “0” AND (iii) the month field is between 1 to 12 AND (iv) day field is between 1 to 31.

The age of death of a profile was defined as the difference between the year of death to the year of birth. We filtered a few thousand entries with negative life span or life span above 100. Those entries are usually due to simple typos such as reporting the year of birth with four digits (e.g “1910”) and the year of death with two digits (e.g. “64”).

#### Comparing Geni to traditional demographic data

The expected life span from the Oeppen and Vaupel study was taken from the male column in their Supplemental Table 1. We used the male column due to slight access of males in our data. The life expectancy data from Geni was on average 1.4 year lower than their study across all years. This finding is quite expected as the Oeppen and Vaupell dataset was ascertained from countries with the highest life expectancy, whereas the Geni life expectancy is from a wide range of countries.

Data from the Human Mortality Database (HMD) was downloaded from France, Belgium, England and Wales, and Denmark for 1820, 1850, 1880, 1910, and 1940. Those countries were selected because they had data for the 19th century and since most of the population in Geni has Western European heritage. The life span probability density function for each country and time point was calculated using the `"dx"` column of the HMD file, which lists the number of deaths in each age. We then averaged the histograms from all four countries, except for 1820, in which we only had data from France (Figure S5).

Let *p*_*t*_ (*x*) and *q*_*t*_ (*x*) be the Geni and the HMD age of death distributions for year *t*, respectively. Define 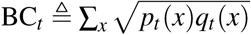, where BC_*t*_ is the Bhattacharyya coefficient for the two distributions for year *t*. The worst Bhattacharyya coefficient reported in the main text was set to BC_*worst*_ = min(BC_1820_, …, BC_1940_).

### Obtaining geographic information

#### Annotating geographic information

In order to analyze the geographical origin of the Geni profiles, we used two sources of information. First, Geni.com has a relatively new feature that automatically translated the geographic location of different events to longitude and latitude. We used this data when it was available.

For the old profiles, we only had free text in the following fields: `birth_location, current_residence, death_location`, and `burial_location` fields of the `JSON` files. This text displayed substantial heterogeneity. In some cases, the genealogists entered the location as part of a short sentence (”Census reports say he was born in Pennsylvania, Cleveland, Ohio, United States”) or in an inconsistent format (”,, New York, USA”). Also some of the text was in foreign languages, such as Russian, French, and Hebrew. To deal with this heterogeneity, we used the geoparsing capabilities of Yahoo! Placemaker service. This web service accepts unstructured text in multiple languages and returns structured location information that includes: the annotated location in a canonical format, the type of the location (e.g. town or state), longitude & latitude coordinates, and the quality of the annotation as a score between 1 (poor) to 10 (excellent). To obtain the most probable location when more than one place could match, we turned on the `autoDisambiguate` option of Yahoo! Placemaker service. In order to process the data efficiently, we constructed a corpus of locations that was comprised of all unique entries in the location fields of the `JSON` files and submitted the unique entries to Yahoo! Placemaker. To increase the reliability of the annotation, we excluded annotations that were too broad (such as continent or country). Next, we filtered annotations with quality level of 8 or above.

#### Validating the annotation accuracy using historical events

We placed profiles in the time-space domain by associating either their birth year with their birth location or their death year with their death/burial location, with the former given a priority over the latter. The year of birth or death was extracted only from fields that had full date information. The years of settlement were taken from the Wikipedia page of each city and reflected the earliest documented Western settlement in the area, which in some cases was before the official incorporation of the city.

### The genetics of longevity

#### Modeling the expected life span

Longevity was defined as the deviation of the age of death from the expected life span based on temporal and environmental factors. As we were interested only in adult death, we restricted the analysis only to profiles that their age of death is above 30.

To calculate the expected life span, we evaluated 10 different models. The following R code describes these models:

~~~
# only gender
model1<- bam(long ~ gender, samfrac=1, data=training_set)

# birth year
model2<- bam(long ~ birth_year, samfrac=1, data=training_set)

# gender and birth year
model3<- bam(long~gender+birth_year, samfrac=1, data=training_set)

# gender, birth year, and birth country
model4<- bam(long~gender+birth_year+birth_country, samfrac=1, data=training_set)

# gender, birth year, temperature (mean and sd)
model5<- bam(long~gender+birth_year+tmp_mean+tmp_sd, samfrac=1, data=training_set)

#gender, birth year, geo location using spline regression
model6<- bam(long ~ gender + birth_year +s(birth_location_latitude, birth_location_longitude, bs=“sos”,k=splines), samfrac=1, data=training_set)

# gender, birth year, birth country, and temperature (mean and sd):
model7<- bam(long ~ gender+ birth_year+ birth_country+ tmp_mean+ tmp_s, samfrac=1,data=training_set)

# gender, birth year, temperature (mean and sd), and geolocation using spline regression:
model8<- bam(long ~ gender + birth_year+ tmp_mean+ tmp_sd+ s(birth_location_latitude, birth_location_longitude, bs=“sos”,k=splines), samfrac=1, data=training_set)

# gender, birth year, birth country, and geolocation using spline regression:
model9<- bam(long ~ gender +birth_year + birth_country + s(birth_location_latitude, birth_location_longitude, bs=“sos”,k=splines), samfrac=1, data=training_set)

# gender, birth year, birth country, and temperature (mean and sd), and geolocation using spline regression:
model10<- bam(long ~ gender+ birth_year+ birth_country+ tmp_mean+ tmp_sd+ s(birth_location_latitude, birth_location_longitude, bs=“sos”, k=splines), samfrac=1, data=training_set)
~~~

Temperature information as GIS was obtained from WorldClim in 2.5min resolution. We averaged the mean temperature reflects the average annual temperature and the standard deviation reflects the per month spread around the mean. We used the birth location of each profile to query the GIS grid and obtain the average temperature and standard deviation. While individuals can migrate from their birth locations, our data show that most migration events are very small especially before 1850. Therefore, we posit that the temperature at place of birth largely reflects the climate that the person experienced.

The country of birth was defined as a factor variable. To increase to associate this covariate, we restricted the data to individuals that were born in the top 25 countries with most profiles, namely: “United States of America”, “Netherlands” ,”Sweden” ,”Germany” ,”United Kingdom” ,”Norway” ,”Estonia” ,”Finland” ,”Canada” ,”Denmark” ,”France” ,”Australia” ,”Belgium” ,”Poland” ,”South Africa” ,”Russia” ,”Italy” ,”Czech Republic” ,”Switzerland” ,”Ireland” ,”New Zealand” ,”Austria” ,”Croatia” ,”Hungary” ,”Spain”.

To find the best prediction for life expectancy, we trained each model with 90% of the profiles and tested the prediction accuracy with the other 10% profiles.

The best model included the gender, year of birth, and the geolocation. All of the covariates of the model were significant. It explained 7% of the longevity of the test set, it had the minimum BIC score, and the minimal mean squared error with the test set.

To calculate the longevity of individuals, we simply subtracted their age of death from the expected life span using this model.

#### Filtering individuals before evaluating genetic models

To further reduce environmental factors, we employed the following filtration steps:

a. Including only individuals with full year of birth and year of death that we can calculate their life expectancy using the model above.
b. Removing individuals that were born after 1910 to avoid ascertainment bias towards early lifespans (Figure S17) or before 1600, which can have lower reliability.
c. Removing deaths prior to age of 30.
d. Removing individuals who died during the American Civil War, WWI, and WWII, which showed a marked increase in death rates of military age individuals.
e. Removing individuals without precise geographical assignment. After finding pairs of relatives (see next section), we filtered additional pairs:
f. We filtered potential twin pairs from the dataset since to avoid erroneous IBD estimation as it was to distinguish between MZ and DZ pairs.
g. We filtered pairs of individuals with an expected IBD over 60% as these are likely to represent genealogy errors.
h. Additionally, we observed that pairs of individuals that were born in the same town had higher death rates within the same year (Figure S8). This pattern was mostly evident in siblings and first cousins but also even in fourth cousins that were born in the same town (but not fourth cousins that were born far away from each other). This pattern is presumably the result of local environmental hazards such as natural disasters or violent activity. We were concerned that these local hazards can confound the analysis as more related individuals tend to live closer, which can induce spurious association between genetic similarity and longevity. To mitigate the effect of such catastrophes, we removed pairs of individuals that died within 10 <u>days</u> from each other. Indeed, filtering these pairs removed the over-representation of deaths within the same year.

#### Heritability with nuclear families

Let *i* be an individual whose father is *f* and mother is *m*. The mid-parent heritability was measured by `lm`, the least-square regression package of `R` to infer the *α*_0_ and *β* of the following model:

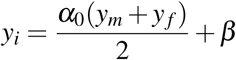

*h*^2^ was set to the estimator of *α*_0_.

For finding a secular trend in the mid-parent heritability over time, the years of birth of the parents were averaged and grouped into bins of 10 years. We employed the mid-parent regression for each bin of data and calculated the maximum likelihood estimate of the heritability and its standard error. Then, we regressed the heritability results on average year of birth of the parents using linear regression with weights that correspond to the inverse of the standard error of each heritability measurement.

The 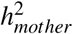 and 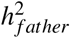 were calculated similarly to the mid-parent design, by inferring *α*_0_ and *β* of the following models: *y*_*i*_ = *α*_0_*y*_*m*_ + *β* and *y*_*i*_ = *α*_0_*y*_*f*_ + *β*, respectively. The reported *h*^2^ was set to 2*α*_0_ in each model.

Testing whether the difference between 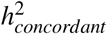 and 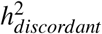 is significant was done by measuring the *p*-value of the interaction term *α*_2_ in the following model using linear regression:

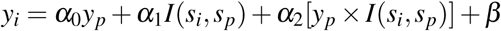

where individual *p* is either the mother or father of *i*, *s*_*i*_ ∈ (male, female) and *I* is an indicator function that returns 1 for *s*_*i*_ = *s*_*p*_ and 0 otherwise.

#### Adjusting relationships

In order to collect and measure the IBD between relatives, we employed the following steps:

a. Finding relative pairs: familial ties can be defined by three basic relationships: A is a parent of B, A is a full sibling of B, and A is an offspring of B. For example, C is the uncle of A, if C is the full sibling of B, and B is the parent of A. For each individual that passed the multiple inclusion criteria, we scanned for all the profiles that are related to him by alternating between these three basic steps to identify siblings, parents, grandparents, great-grandparents, uncles, cousins or cousins once removed up to 4th cousins.
b. Maximizing relationships: in the presence of consanguinity, the genealogical relationships are not unique since a relative can be reached by multiple paths. To address this issue, we assigned the closest possible relationship if more than one path was detected. For example, if A and B could be second cousins or fifth cousins, we assigned them to the second cousin group.
c. Calculating identity coefficients: another complexity of consanguinity is that the expected IBD between related individuals can be higher than their genealogical relationships (Figure S9). In addition to assigning pairs of relatives to genealogical classes, we also calculated the expected IBD taking into consideration all potential paths using relatives up to nine generations. For this task, we employed IdCoeff [1], which calculates Jacquard’s 9 Condensed Coefficients of Identity. These coefficients are an extension of the IBD probabilities and represent all possible configurations of a bi-allelic autosomal site between a pair of individuals, given that the parent of origin does not matter (see inline figure on the right). Since IdCoeff was designed for relatively small pedigrees, we modified the source code to accept topological sorted pedigrees, which removed the *O*(*n*^2^) pre-processing time of registering the pedigree in the computer memory to *O*(*n*log(*n*)) (code is available from the authors). Running the modified IdCoeff on all possible genealogical pairs of relatives took approximately 25000 hours of CPU time (a month of a server with 15 parallel processes).
d. Calculating the expected IBD: Let *r*_*i*__*j*_ be the IBD probability between individuals *i* and *j* in the presence of consanguineous marriages. Then (see [2]):

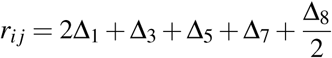

where ∆_*i*_ is the *i*-th identity coefficient for the *i, j* pair according to the IdCoeff output.

**Figure:**
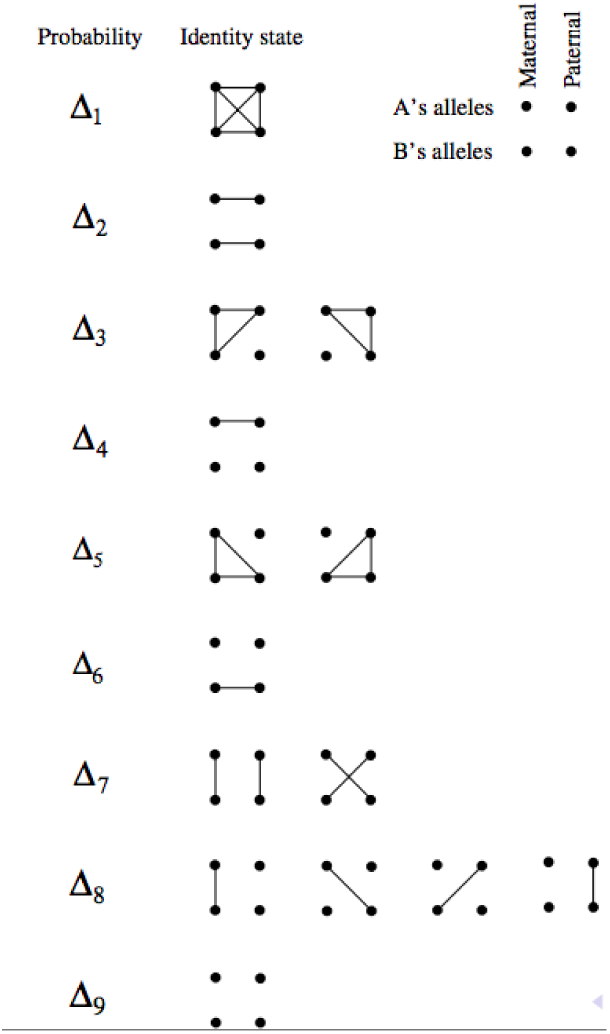
Nine possible identity states

#### Measuring dominance variance

Throughout this manuscript, the phenotypic correlation is defined as:

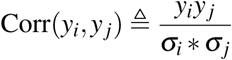

where *y*_*i*_, *y*_*j*_ are the longevity of individual *i* and *j*, respectively, and *σ*_*x*_ is the standard deviation of all individuals in class *x*.

The ∆_7_ allelic configuration mediates the variance of *v*_*d*_ and mainly appears in full sibs (Figure S10). To estimate this factor, we sought to compare full-sibs to parent-child pairs, in which ∆_7_ *≈* 0. This comparison is immune to confounders due to additive and epistatic factors as the average IBDs of both relative groups were the same with 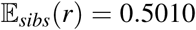 and 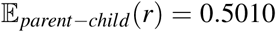, and therefore, should mediate on average similar additive and epistatic vavriance. To further reduce potential biases due to sex differences, we limited the analysis only to males, resulting in 157, 600 father-son pairs and 159, 000 brother pairs.

However, one potential caveat in this setting is that brothers can be more correlated due to transient environmental effects, such as local catastrophes that are unaccounted in our model. To illustrate this, consider a family of a father and two sons of ages 70, 35, 30 years old, respectively that live in the same town. In the case of an extrinsic catastrophe that kills all of these individuals, the two siblings will be more correlated than the father-son pairs. Importantly, the correlation difference simply reflects the fact that the two sibs were born in similar years compared to their father and has nothing to do with dominance. This excess in correlation is not unique to our data and was documented in previous studies. For example, a classic paper by Rao et al. [11] inspected the correlation of height and weight in nuclear families. Despite adjustments of the phenotype to the age of the child, they found a strong decline in correlation as a function of the difference in year of birth between siblings.

To address that, we tested the two models using non negative least square regression based on the `R` package `nnls`:

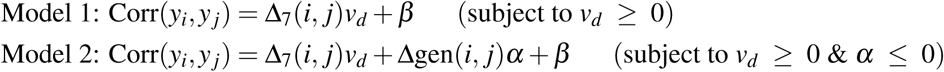

where ∆gen is the absolute difference in the year of birth of the two profiles, which can reflect changes in the environment over time. The first model estimated *v*_*d*_ = 0.24. The second model estimated *v*_*d*_ = 4.0 and *α* = −0.0018.

Several pieces of evidence support the second model: first, the BIC(model1) *−* BIC(model2) = 41, suggesting that Model 2 is more appropriate. Second, we also regressed:

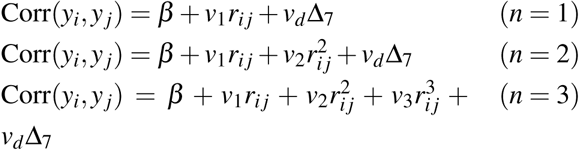

but this time only with pairs of brothers. Again, we found that *α* = *−*0.0015 and was statistically different than 0 (*p <* 0.0005). This shows that ∆gen explains similar phenotypic correlation to Model 2 while *v*_*d*_ is nearly fixed. Third, Model 1 estimated *v*_*d*_ = 0.24 *±* 0.03. This implies that the entire correlation observed in the MZ twins (0.24) is attributed to dominance variance. However, the heritability of sex-concordant parent-offspring pairs was estimated to *h*^*2*^_*concordant*_ = 0.21 *±* 0.01. This estimator is robust to dominance variance, suggesting that *v*_*d*_ was largely overestimated by Model 1. Therefore, we estimate the that the dominance variance of longevity is around 4%.

#### Model fitting

We evaluated the following three nested models:

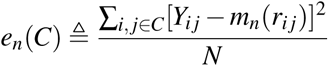

Fourth cousins, which display almost zero IBD probabilities, showed longevity correlation of 2.0% that was significantly different than 0 (*p <* 10*−*^24^). This correlation might be due to the fact that fourth cousins live much closer to each other (median distance: 75km) than complete random pairs of individuals in our data (median: ~2000km). *β* was included in the models to allow positive longevity correlation in these far related individuals.

The models were fitted to the data using non negative least squares with the `R` package `nnls`, forcing *β, v*_1_*, …, v*_3_ *≥* 0. The value for *v*_*d*_ was set to 4.0 according to the results above. For testing whether the model was statistically significant, we used nested ANOVA. Both this function and the BIC calculation were done using custom functions in `R`. For cross-validation, we removed each class of relatives in Table S2 and fitted the three models with the remaining data. Then, we calculated the mean squared error based for the *C* class by:

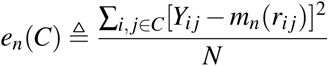

where *Y*_*i*__*j*_ is the observed longevity correlation for the *i, j* pair that is part of the *C* class. *m*_*n*_(*r*_*i*_) denotes the longevity correlation prediction of the *n* model (e.g. n=1) for the *i, j* pair based on their IBD readout, which is equal to *r*_*i*__*j*_. *N* is the number of pairs in the class. The total MSE of the k-th model was set to 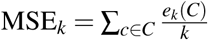.

For consistency with the heritability of dominancy, we also repeated the regression of the three configurations with ∆gen (difference in age of birth) as an additional covariate. This procedure had no effect on our conclusion. The epistatic terms were not significantly different than zero, did not improve the MSE, and still showed a large difference in their BIC values compared to the additive model.

The Danish twin pairs underwent the similar adjustment and pre-processing steps as the Geni data. Namely, we converted their age of death to longevity using the same model that takes into account sex, year of birth, and the geolocation of birth. As we did not know the exact geo-location, we set the birth place to Copenhagen, the most populated metropolitan in Denmark. We note that due to the small size of this country (without Greenland) assignments to other coordinates with Denmark has virtually no effects on the life expectancy estimation of our model. In addition, similar to the Geni relatives, we removed MZ-twins that died before age of 30 or during WWI or WWII. Finally, we adjusted the twin correlation to reflect the measured *v*_*d*_.

The only difference from the Geni processing steps was that in the absence of access to the exact dates of death, we could not filter individuals that died within 10 days of each other. This caveat could create a slight increase in the longevity correlation of MZ twins compared to other pairs in our. Such bias would be in favor for epistatic models and therefore does not affect our conclusion about the role of additivity.

MZ perdiction was measured by evaluating the *n* model in *r* = 1.

We note that we deliberately decided to use Haseman-Elston (HE) linear regression framework rather than a variant component method (linear mixed model). The main issue is that we needed to exclude certain types of relatives to reduce correlation due to environmental catastrophes (such as fin fig. S8). Removing relatives is not adequate for the VC method as it creates a hole in the relationship matrix and the current theoretical framework does not allow this type of data cleaning. In addition, recent quantitative studies showed that HE regression provides very similar results to methods that operate on the kinship matrix and we do not expect to get much different results with this method [4, 6]. To address the correlated data points in our analysis, all standard errors of the model estimators are based on a bootstrapping technique using 100 iterations.

#### Addressing potential environmental confounders and more complex models

1. Adjusting household effects The correlation of longevity between relatives can be induced to shared household effect such as access to the health care system, socio-economic status, or dietary habits. To mitigate these effects, we adjusted the life expectancy of an individual using data from their spouse similar to [7]. The idea is that if the spouses are not related to the individual of an interest, than the correlation of longevity in couples includes shared household effects. To best fit the data, we extended our life expectancy model and considered the following configurations in R: ~~~
# sex, birth year, and the spouse life span:
model_s1 <-bam(long1~ birth_year + sex1+long2, samfrac=1, data=training_set)

# sex, birth year, the birth geolocation (with splines), and the spouse excess longevity
model_s2 <-bam(long1 ~ birth_year + sex1+ s(lat1,lon1,bs=“sos”,k=1000) + ex_s, samfrac=1, data=training_set)

# sex, birth year, the birth geolocation (with splines), and the spouse life span
model_s3 <-bam(long1 ~ birth_year+ sex1+ s(lat1,lon1,bs=“sos”, k=1000)+ long2, samfrac=1, data=training_set)

# sex, birth year, birth geolocation (this is the previous model and it was evaluated as a control)
model_s4 <-bam(long1 ~ birth_year+ sex1+ s(lat1,lon1,bs=“sos”,k=1000), samfrac=1, data=training_set)

# the sex, birth year, and spouse life span:
model_s5 <- bam( long1 ~ birth_year+ sex1+ ex_s, samfrac=1, data=training_set)
~~~ The data consisted only of couples that are not genetically related according to our tree data to avoid confounding the household correlation with genetic correlation. In addition, we only considered couples that both individuals dies after age of 30. Finally, if the individual remarried to other person, we averaged the life expectancy of their spouses before calculating the model. The parameters were inferred using 90% of the data (approximately 2.5 million data points) and the evaluating of the model was done with the other 10%. We found that the model s3 gave the best results S11. It has the lowest BIC value, the lowest mean squared error in the test set, and explained the highest fraction of the longevity variance. In addition, all of the covariates were statistically significant according to ANOVA (*p <* 0.01). In overall, spouse longevity helps to explain an additional 1% (s.e. = 0.2%) of the longevity variance compared to the original model that just considered sex, year of birth, and geolocation. These results are consistent with previous longevity studies in twins [9] and nuclear families [7] that also estimated a similar contribution of the shared sizable household effect. The decomposition of genetic variance of the spouse-adjusted longevity (n=450,000) had no effect on our conclusions. Additivity explained *h*^2^_concordant/relatives_ = 16.0% (s.e.= 2.2%) of the variance, dominancy explained 3% (s.e. = 5.9%), and the epistatic values converged to zero (quadratic s.e. = 0.1%, cubic s.e = 7.4%). Again, BIC and RSE estimates disfavored the epistatic terms. We did not compare this condition to the Danish MZ twin dataset since we could not apply the same spousal correction for the Danish MZ twins. The only difference was that the new model estimated a lower dominance variance *v*_*d*_ = 2.3% to longevity based on the difference in the longevity correlation in sib pairs versus parent-offspring. The lower observed dominance is not surprising. In the the current analysis, the father’s phenotype includes an adjustment based on the longevity of his wife, which shares 50% of her genome with the offspring. Thus, the parent-offspring correlation is expected to be higher. Since dominance variance is calculated as the difference between the correlation of sibling minus the correlation of parent-offspring, the outcome is expected to be lower. In any case, we also evaluated the three nested models (additivity, epistasis of two genes, and epistasis of three genes) after adjusting the correlation to 4% dominance and using the spouse-adjusted phenotypes. The results did not change and the epistatic terms converged to zero. We sought to measure the shared household environment from a different angle. To this end, we analyzed the longevity correlation in (a) pairs of same-sex individuals whose respective spouses are siblings and (b) pairs of same-sex individuals whose spouses are first cousins. For simplicity, we will dub the former pairs as Class I and the latter pairs as Class II. For example, consider that Alice married to Bob, and that Bob and Casey are brothers, and that Casey and Dora are married. Alice and Dora represent class I pairs (pair of individuals whose spouses are sibling). Similarly if Bob and Casey are first cousins, then Alica and Dora are case II pairs (pair of individuals whose spouses are first cousins). We filtered the pairs according to the previous steps (no deaths during major wars, only sex-concordant pairs, etc). We also filtered any pairs that are also related (up to fourth cousins). For example, if Alice and Dora happen to be third cousins, we removed them from the analysis. We were concerned that Class II could show lower correlation due to larger geographic distances of first cousins. To address this issue, we measured the distance in the place of birth of the kids of each of these pairs. For example, if Alice gave birth to Adam and Dora gave birth to Diane, we measured the distance of the place of birth of Adam and Diane. For that two classes, we only retained pairs whose kids were born at the same place to reduce. Our hypothesis was that child birth better represents the environment of individual later in life and therefore would help to capture additional environmental correlations on top of our adjustment to the place of birth for life expectancy. After controlling for these these potential environmental correlations, the extra correlation of Class I over Class II can represent:
  a. Correlations due to similar household environment
  b. Correlations due to assortative mating patterns. For example, if Bob is tall then it is likely that his brother Casey is also tall since the heritability of height is 80%. Due to assortative mating, it implies that both Alice and Dora are also likely to be taller than average. However, since the correlation of height in cousins is likely to be smaller, then we expect that correlation due to assortative mating with respect to height should be smaller in cousins. Similarly, if there are certain factors of longevity that play a role in assortative mating, they are likely to be more correlated in Class I pairs than Class II pairs.
  c. Correlations due to grieving effects. Previous research have reported an increased likelihood of a person to die after the death of their spouse [5] or the their sibling [12].For example, if Bob and Casey are brothers, the death of Bob can affect Alice and Casey, which could eventually affect Dora. We hypothesize that these effects are less likely between first cousins. Therefore, they should contribute to the excess of correlation in Class I over Class II.
  Taken together, the difference between the longevity correlation of Class I pairs and Class II represents the net effect of the three conditions above. Since these conditions are likely to be non-negative numbers, the difference between the class is likely to reflect an upper bound of the shared household environment. Measuring the difference between Class I and Class I, we found that Class I had only 1.5% additional correlation in their longevity, which is less than tenth of the measured heritability. This likely upper bound argues against strong shared household effects in longevity. It is also important to note that household effects are more likely to inflate the correlation of close relatives in our model, such as sibs and half-sibs. As such, they can create spurious epistasis signal rather than inflating the additive signal. Therefore, our results regrading absence of epistasis are unlikely to be affected by if unaccounted household effects still exist.
2. **Only female lines** Our results with the Y chromosome and mitochondria data showed that our pedigree has a low percentage of errors. We wondered whether our results are affected by these errors. To address this issue, we focused on relatives that are purely due to shared maternal lines, because these edges in our graph are much more reliable (0.3% error rate versus 1.9%). We scanned all pairs of relatives and find the shortest genealogical path between them. Next, we filtered any pair of relatives that the intermediate people along their genealogical path did not include females. This process resulted with over 300, 000 pairs of relatives. Next, we fitted the three models to the data. Again, the results were highly similar to the full model that included pairs of relatives due to male lines. Additivity explained 16.9% (s.e.= 2.2%), the epistatic values converged to zero (quadratic s.e.= 0.5%, cubic s.e=6%), and all model diagnostic strongly disfavored any epistatic interactions.
3. **No inbred lines** Abney et al. [2] showed that a population that is the product of consanguineous marriages contains additional components of variance that can contribute to the phenotypic correlation: *v*_*h*_, Corr_*h*_(*a, d*), and 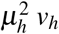 denotes the variation component due to sharing two pairs of identical alleles between a pair of relatives. Corr_*h*_(*a, d*) denotes the correlation of the additive and dominant effects. 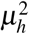 denotes the inbreeding depression in homozygous individuals. The phenotypic correlation of a relative pair due to dominancy in an inbred population is:

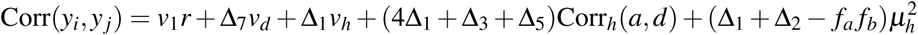

where *f*_*a*_ and *f*_*b*_ denote the inbreeding coefficient of each individual in the pair. While the average values for ∆_1_ to ∆_6_ are extremely small in our pedigree (< 0.1%), we sought to evaluate epistatic interactions in the absence of such confounders. For that, we retained only pairs of individuals whose ∆_1_ + … ∆_6_ = 0. With this process, we still had over 2.6million pairs of relatives to evaluate our models. We did not find any difference in the results between this outbred data points and the model with the 3.2million relatives that included also inbred relatives. Additivity explained 15.8% (s.e.=4.2%) and the epistatic terms converged to zero (quadratic s.e.= 0.1%, cubic s.e=0.1%).

### Analyzing familial dispersion

For all migration events, we only used data from individuals that we had an exact date of birth AND high quality birth location as defined above. All of the results below are available as a single R script from the authors.

#### Measuring migration distance

The migration distance corresponds to the great circle distance between the birth location of each pair of profiles. We transformed the distance to log scale according to log_10_(1 + *x*), where *x* is the distance in km. We only analyzed pairs with high quality birth location as defined above and exact date of birth. The year of birth was averaged between the pair of individuals. The migration distance of males was defined as the distance between the birth locations of father-offspring pairs. The migration distance of females was defined as the distance between the birth locations of mother-offspring pairs. The marital radius was defined as the distance as the distance between the birth locations of spouses. The plots were smoothed using a rolling average with a window of ten years. The raw data before smoothing can be seen in Fig. S14 - Fig. S13.

Profiles from Europe were defined as profiles that were born in the following country codes (according to current political borders): ‘AL’, ‘AT’, ‘BA’, ‘BE’, ‘BG’, ‘CH’, ‘CY’, ‘CZ’, ‘DE’, ‘DK’, ‘EE’, ‘ES’, ‘FI’, ‘FO’, ‘FR’, ‘GB’, ‘GI’, ‘GR’, ‘HR’, ‘HU’, ‘IE’, ‘IS’, ‘IT’, ‘LT’, ‘LU’, ‘LV’, ‘MC’, ‘MK’, ‘NL’, ‘NO’, ‘PO’, ‘PT’, ‘RO’, ‘SE’, ‘SI’, ‘SK’, ‘SM’, ‘VA’.

Profiles from North America were defined as profiles that were born in the following country codes (according to current political borders): ‘US’, ‘CA’.

#### Measuring identity by descent between couples

To measuring identify by descent, we employed the procedure in section “Adjusting relationships” to individuals that were married to each other. We only included individuals that are non-founders. The plot was smoothed using a rolling average of two years. To test isolation by distance, we regressed the average identity by descent per year on the martial

#### Azimuth

Distances on longitude and latitude was calculated using the procedure described by [10]. We tested whether the average migration on WE axis was longer than the NS axis. We also tested whether the number of migration events with longer NS distance are more prevalent than WE migration versus NS migration.

#### Fusing datasets with FamiLinx

FamiLinx includes the profile-id of each person in our database (without the names). Our data use agreement allows not-for-profit researchers to consent participants in order to obtain their Geni profile-id to identify them in our the FamiLinx data. Such usage is conceptually similar to the collaborative nature of Geni.com. However, it is important to note that we strictly prohibit any re-identification without the consent of the participant.

To simplify the collection of the Geni profile-id, we constructed a simple web button that researchers can integrate into their website. The button is based on client side Java script and uses the Geni SDK. If the user consent to contributing their Geni profile, they can click on the button. This sends a signal to the Geni.com server and creates a pop-up for the user that asks for their Geni username and password over secure HTPPS communication. Users without a Geni profile can create a new profile as part of this process. Importantly, this pop-up window is served directly from Geni and the researcher cannot see this transaction. Next, the user’s client receives a Json message from Geni that includes the profile-id and some basic account information. Our client side script transmits this message to the server (the researcher), which can now register this information along with other information form the user (e.g. genome, phenotypes, etc…). Then, the researcher can search the FamiLinx tree for the user profile-id and overly the phenotypic or genomic information with the tree structure.

Fig. S18 shows the sequence of events and the experience of the user. We tested and validated this button in DNA.Land and were able to collect profile-ids of over one thousand individuals using this process. Researchers that are interested to test this method can visit:

<u>https://teamerlich.org/geni-integration-example/</u>. The code is available on: <u>https://github.com/TeamErlich/geni-integration-example</u>.

## Supplementary Figures

**Figure S1:**
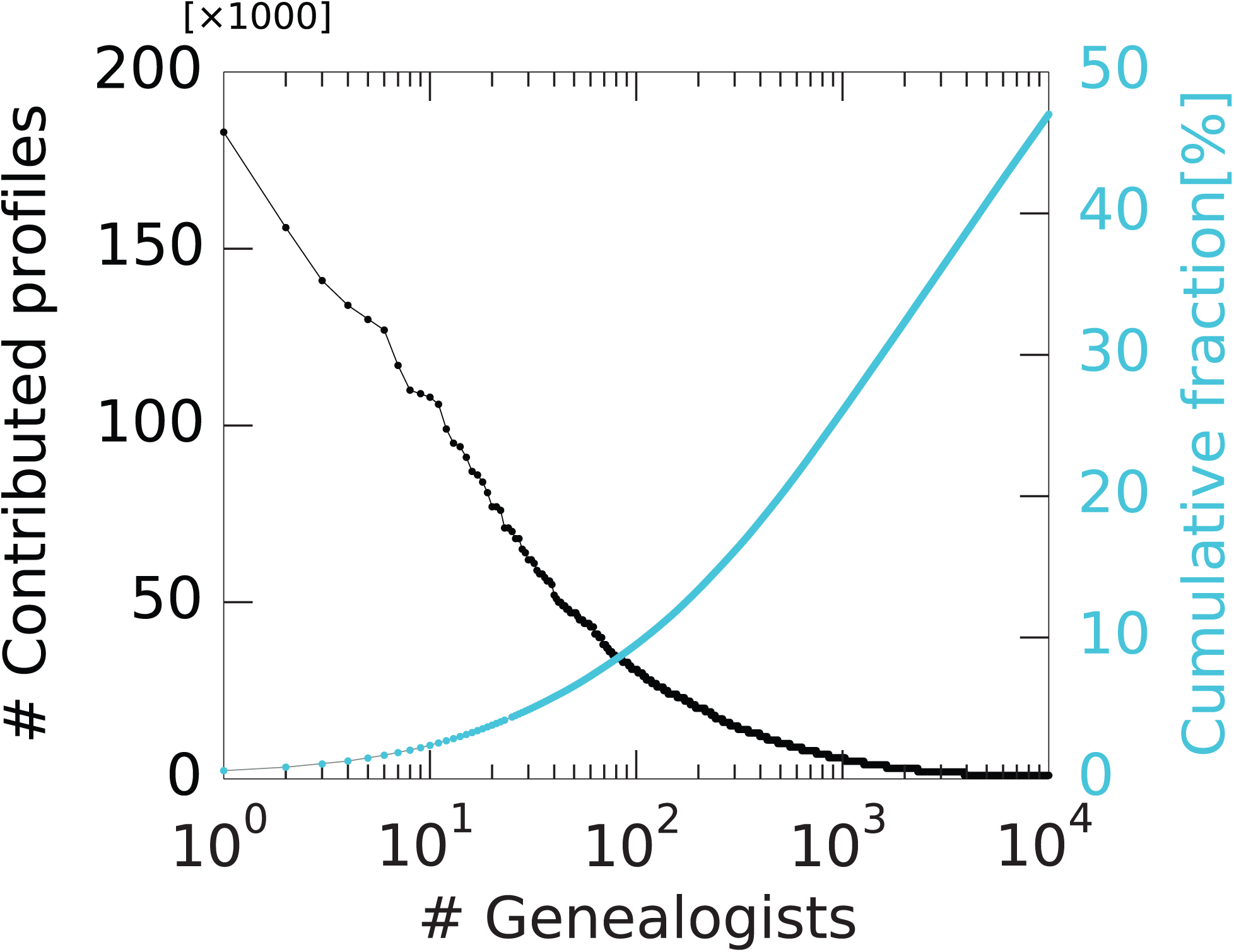
The inferred contribution of profiles by the top ten thousand genealogists (black: number of profiles contributed by each genealogist sorted based on their contribution; light blue: the cumulative distribution of profiles with a known genealogist)

**Figure S2:**
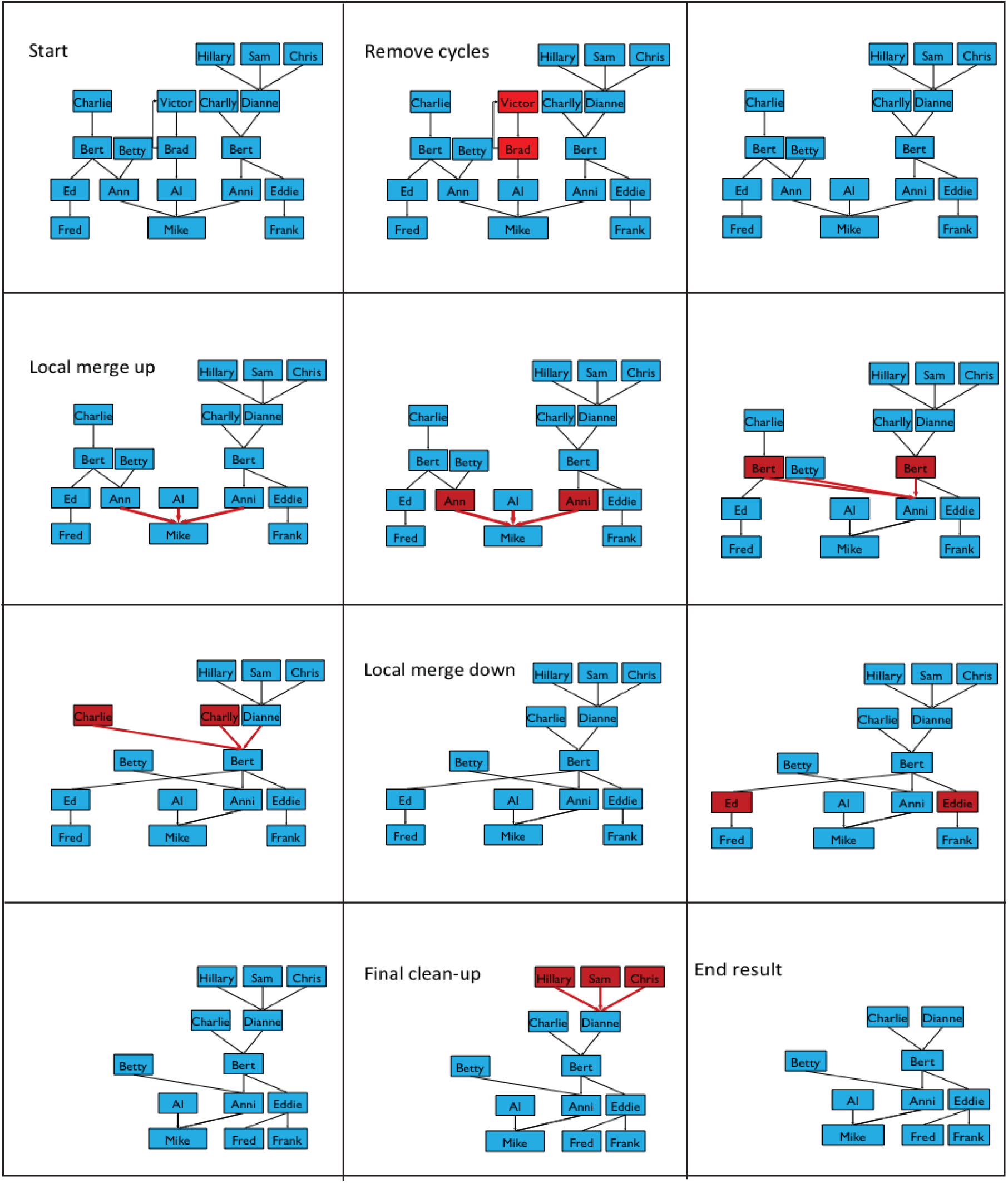
An illustration of the steps taken to clean the Geni graph. Union nodes are not shown for clarity.

**Figure S3:**
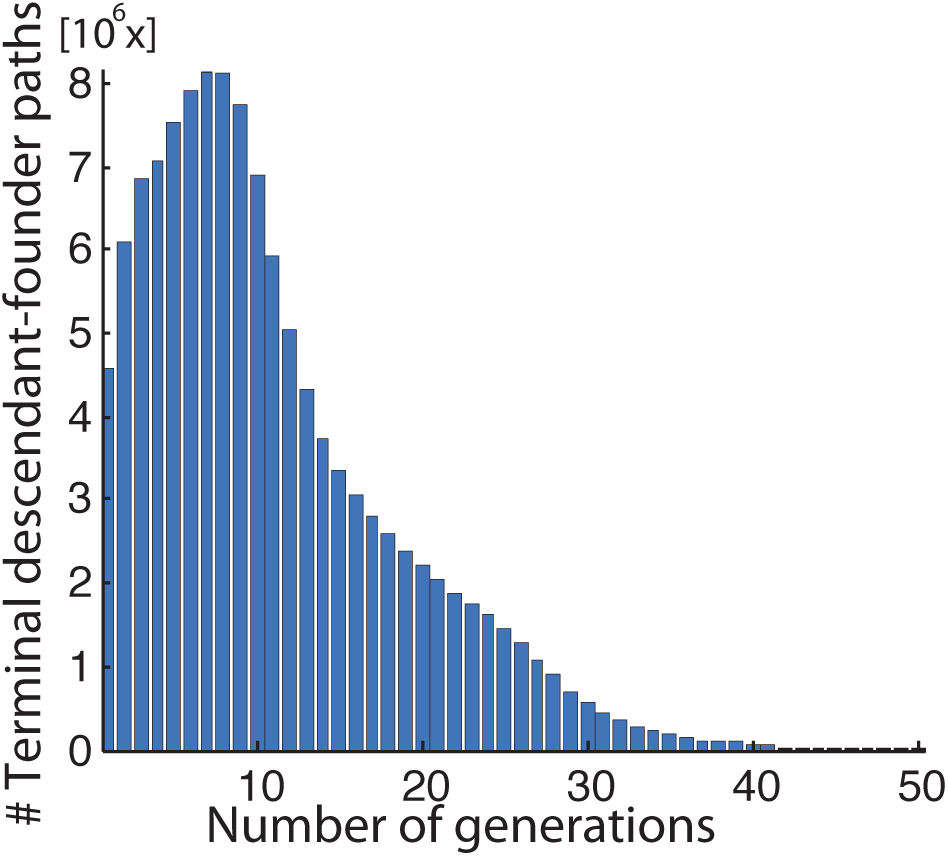
The distribution of number of generations between terminal descendants and founders.

**Figure S4:**
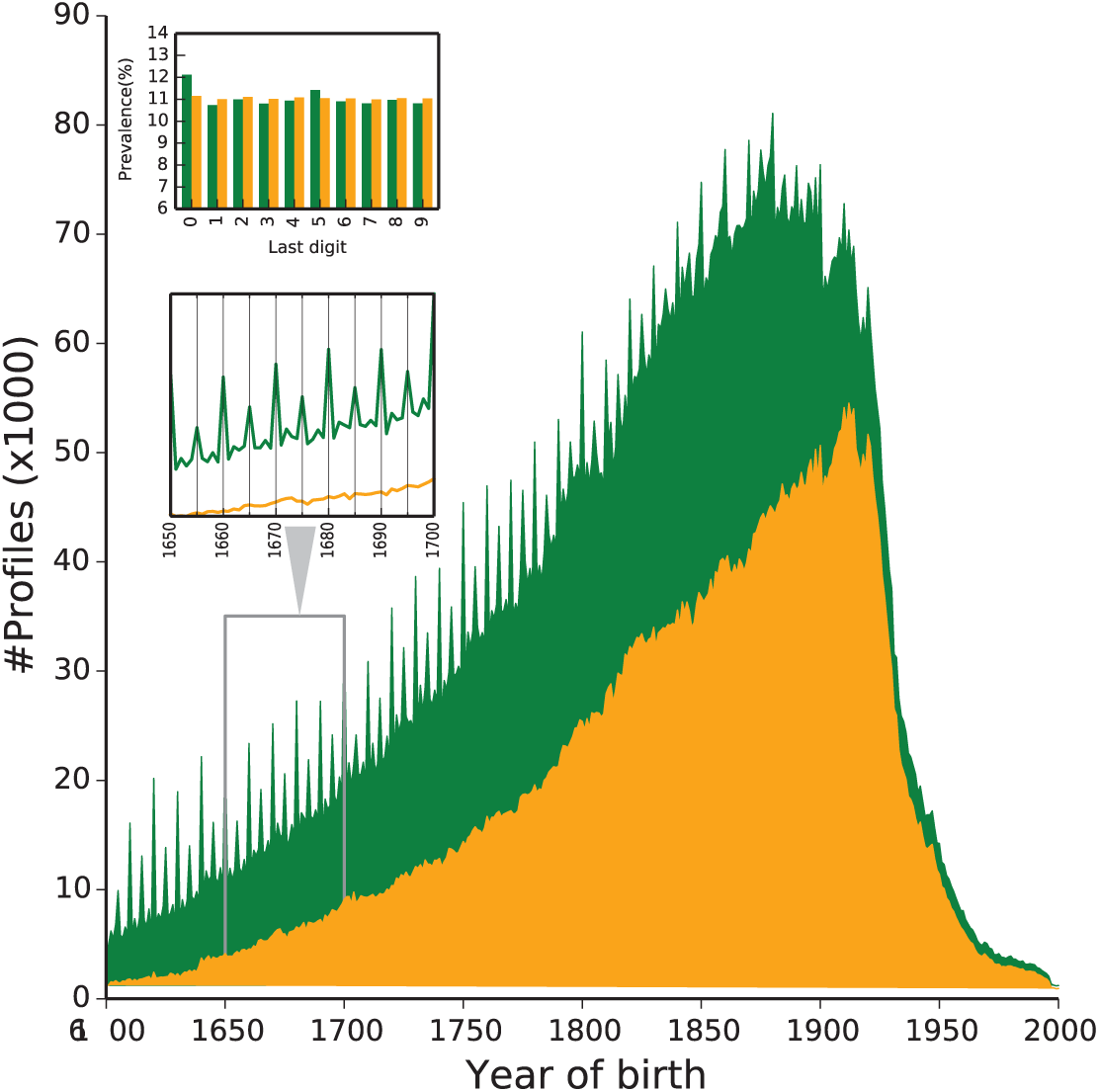
Distribution of the profiles year of birth since the 16th century. Without filtering (green), round decades are overrepresented, suggesting imprecise data. Retaining profiles with exact dates (yellow) removes this pattern.

**Figure S5:**
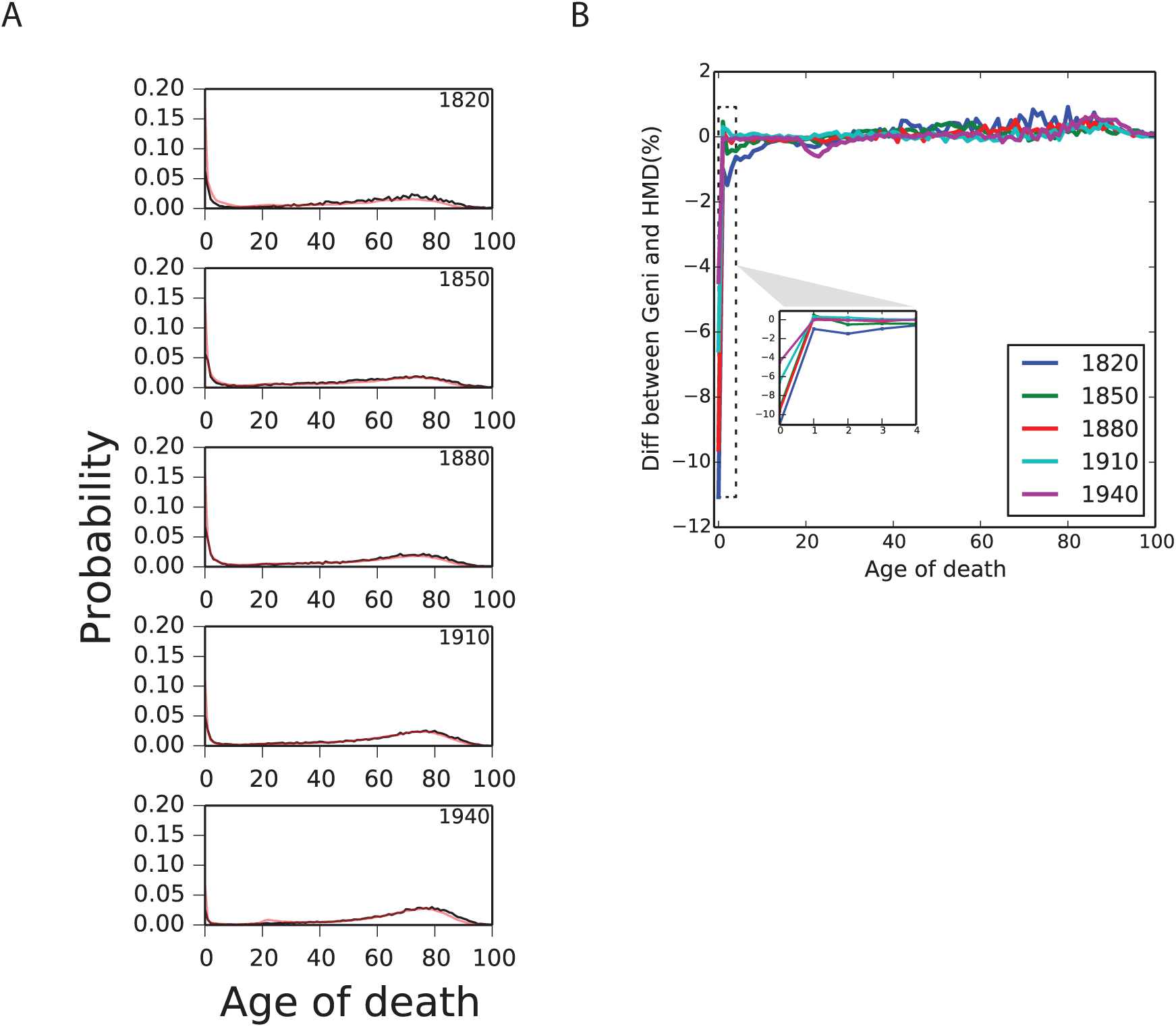
Comparing Geni to HMD (A) The age of death probability density function (PDF) in Geni (black) versus HMD (red) for 1820-1940 (B) The difference between the HMD and Geni data. Each curve represents the subtraction of the HMD’s age of death PDF from the Geni one. The only systematic difference is underestimation of first year of birth deaths in Geni.

**Figure S6:**
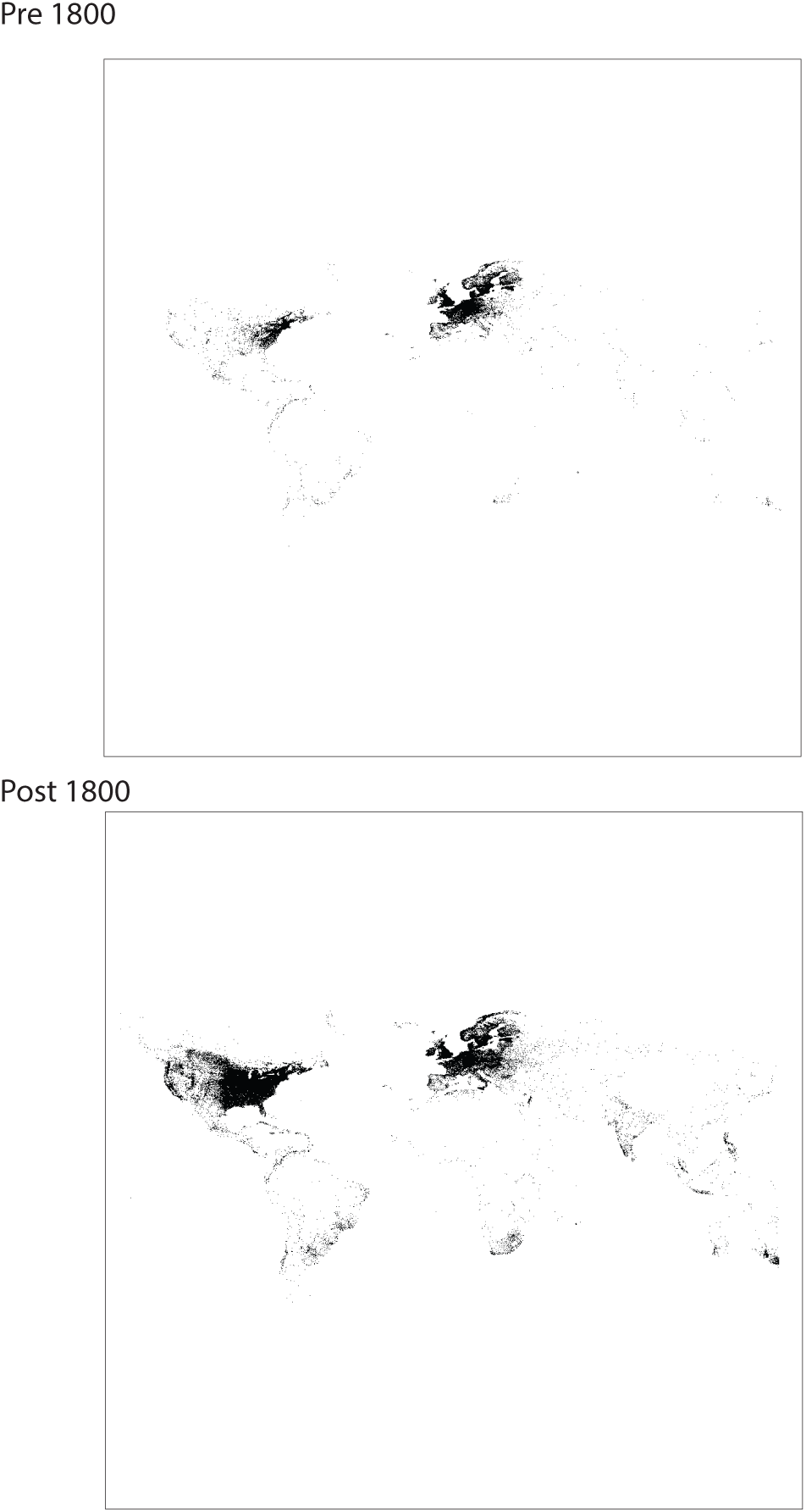
Geographic distribution of Geni profiles pre and post 1800.

**Figure S7:**
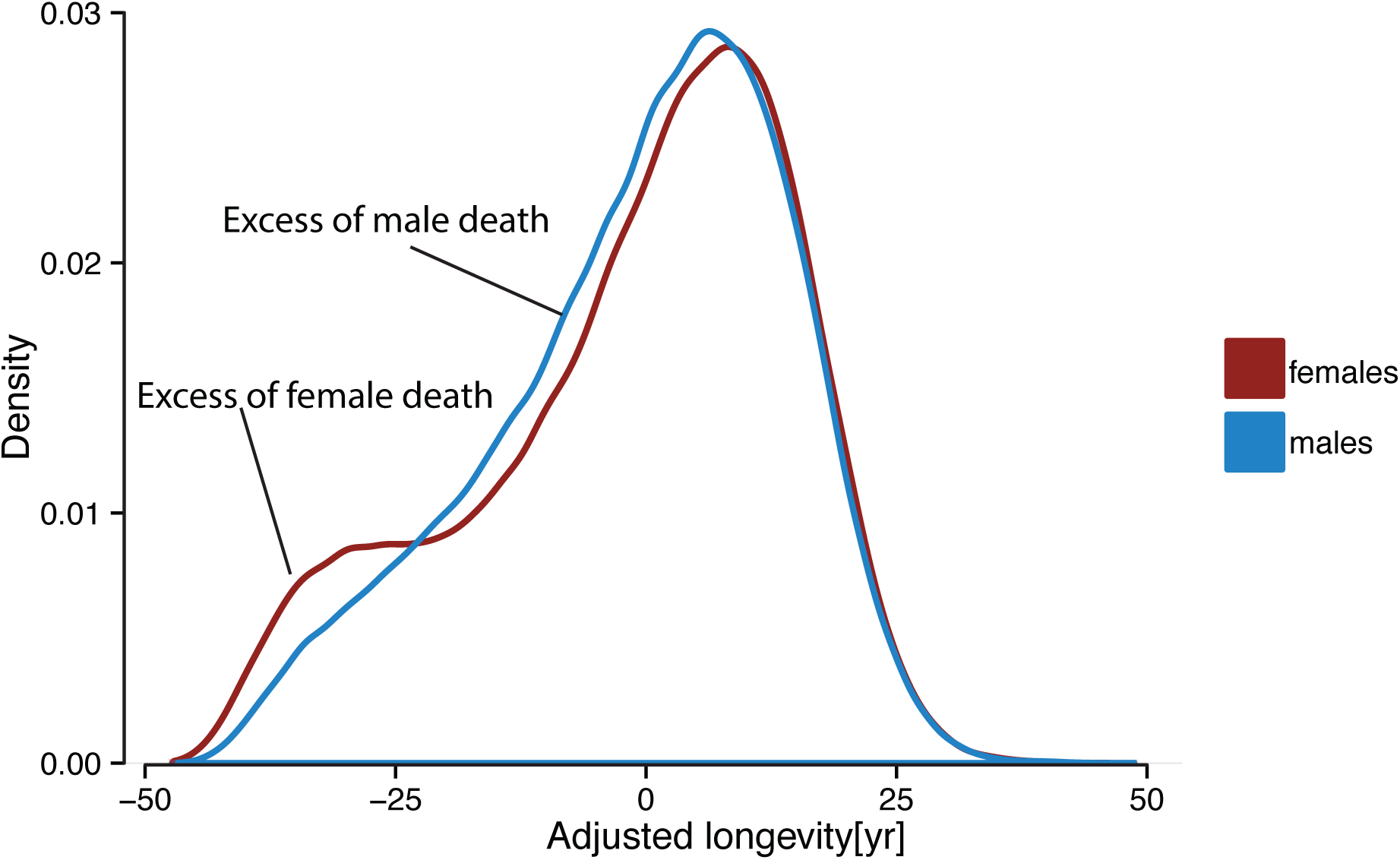
The distribution of the adjusted longevity in all children considered for the mid-parent design. The distribution shows excess of very early female death around child bearing ages whereas males show higher rates of death after those years.

**Figure S8:**
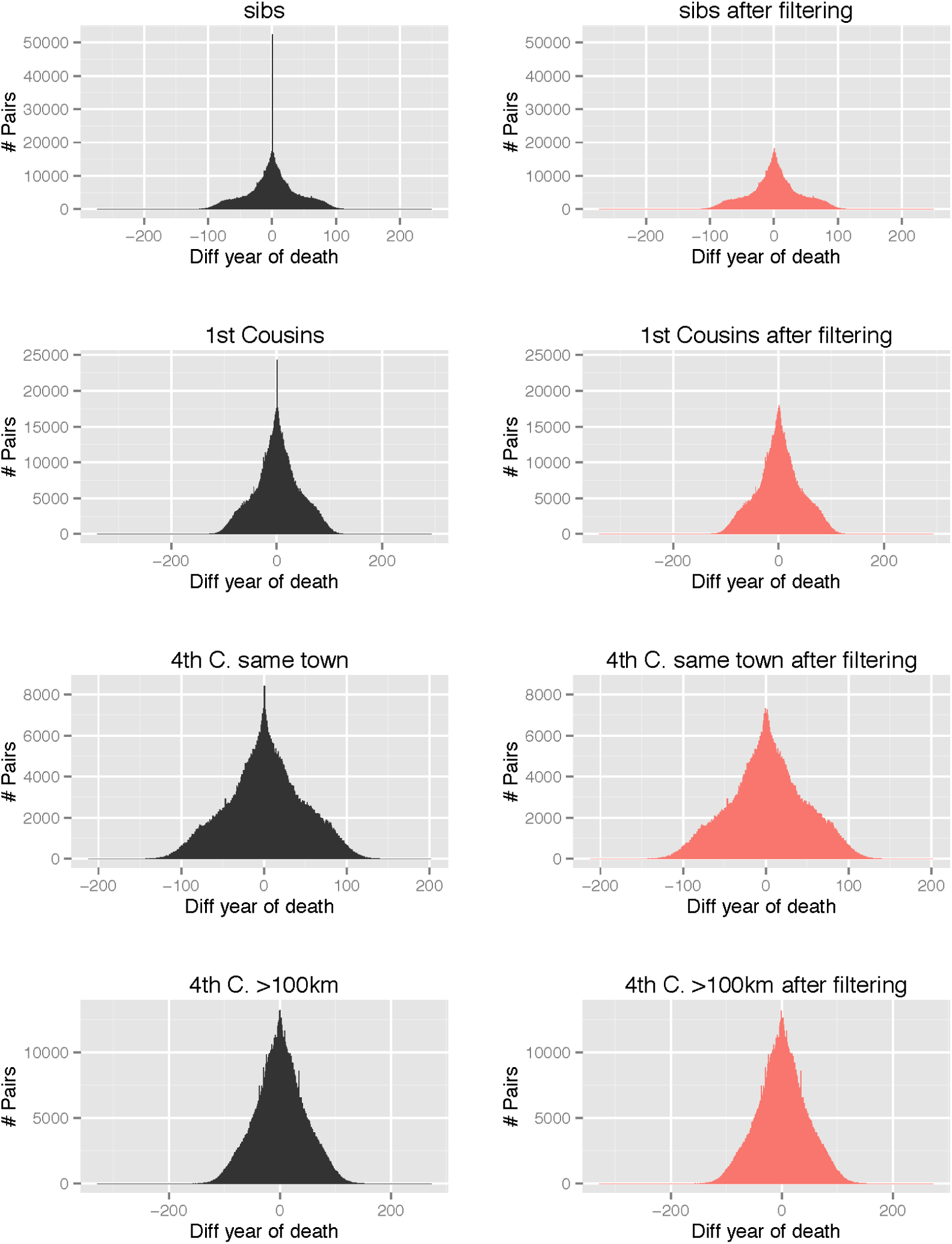
The effect of potential environmental hazards. The histograms present the difference in year of death between various types of pairs of relatives (from top to bottom: siblings, first cousins, 4th cousins that were born in the same town, 4th cousins that were born more than 100km from each other). Left (black): histograms before filtering. Higher death rates within the same year (arrow) are evident in all cases of relatives that were born in the same town. Right (light red): same data after filtering pairs of relatives that died within 10 days. The over-representation is removed, suggesting that this effect is attributed to abrupt environmental hazards such as natural disasters.

**Figure S9:**
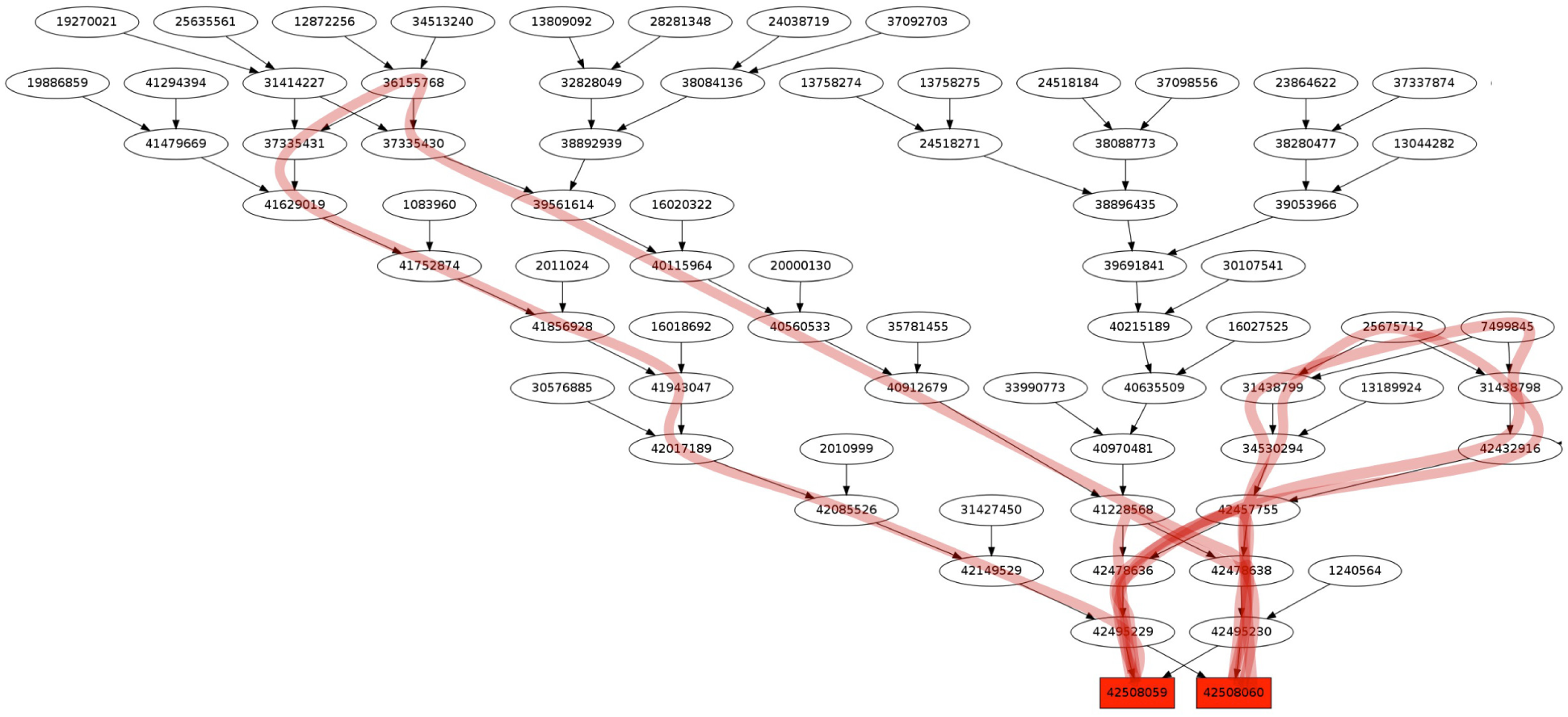
An example of two siblings that are the product of multiple relative marriages. Red - additional paths due to consanguinity. The expected IBD of this pair is 0.565 instead of 0.5. For de-identification, the Node IDs represent the internal representation in our analysis script and not the actual profile ID in Geni.

**Figure S10:**
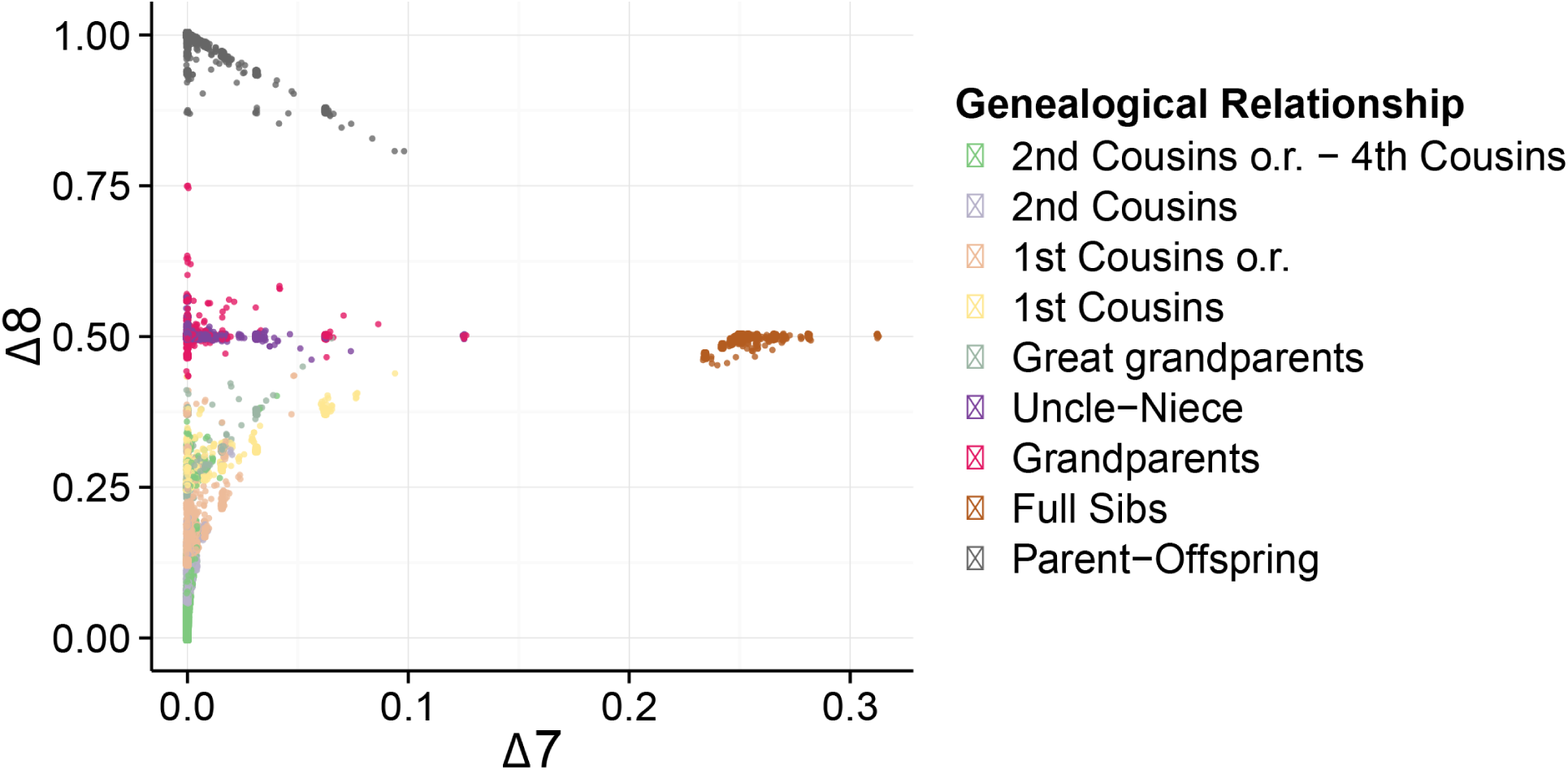
The distribution of ∆_7_ vs. ∆_8_ in our data. Notice that only full sibs exhibit high frequency of the ∆_7_ configuration.

**Figure S11:**
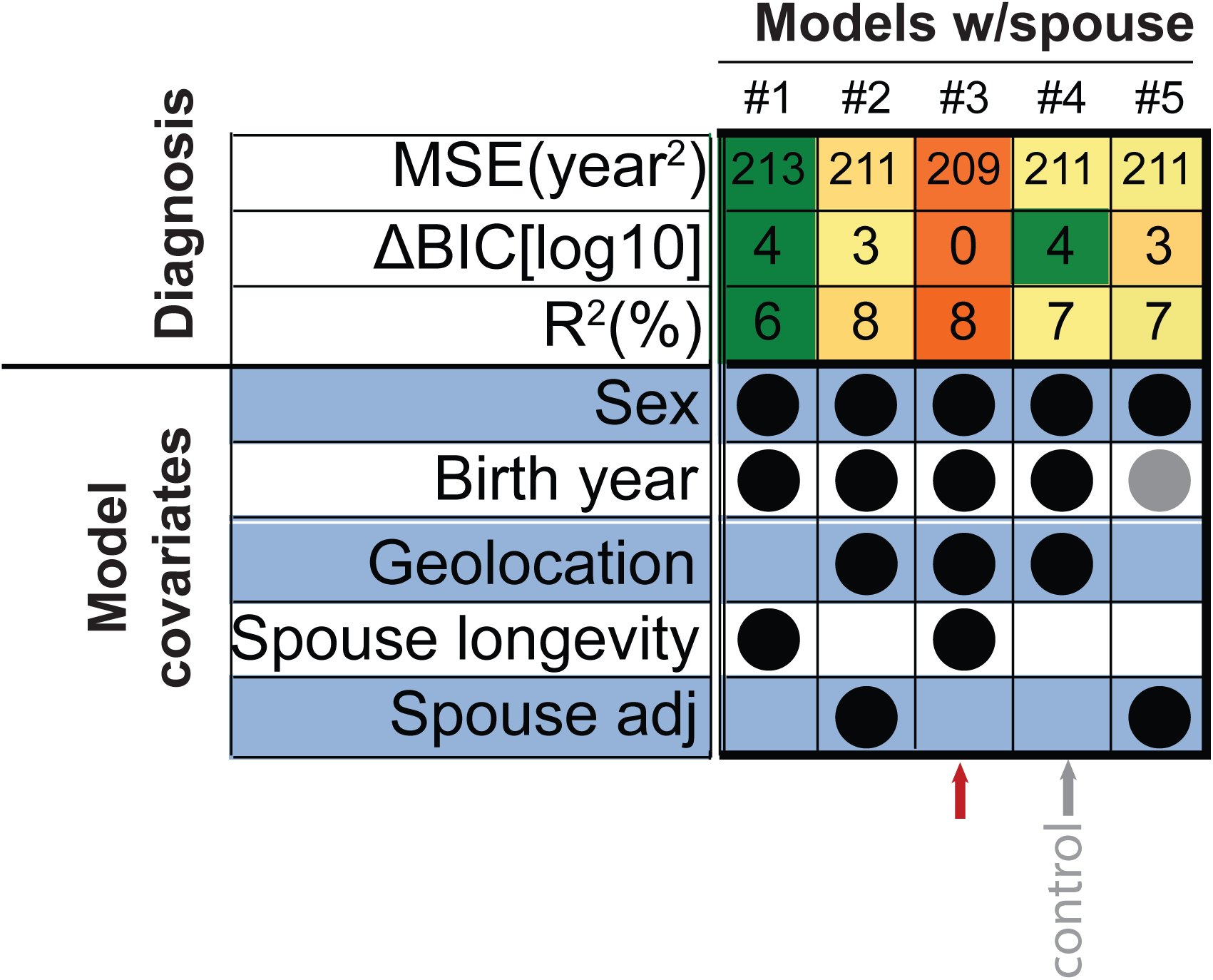
Adjustment of longevity while adding information from spouses of individuals. MSE is the mean squared error per individual; ∆BIC is the difference of the Bayesian Information Criterion from the best model after log_10_(*x* + 1) transformation. Sex is a two level factor (male/female). Geolocation is the longitude/latitude location of birth and was modeled using splines on spheres. Spouse longevity is the age of death of the spouse. Spouse adj. is the longevity of the spouse based after adjustment using model #4 of the main text. Closed circles represent covariates in each model. Gray/Black: statistically in/significant (*p <* 0.01) covariate. Orange to green: most desired to least desired diagnosis outcome. The control model (#4, no spouse information) as a reference point. The best model is #3, which adjusts longevity based on sex, year of birth, geolocation, and spouse longevity. This model had the lowest MSE, smallest BIC, and best *R*^2^.

**Figure S12:**
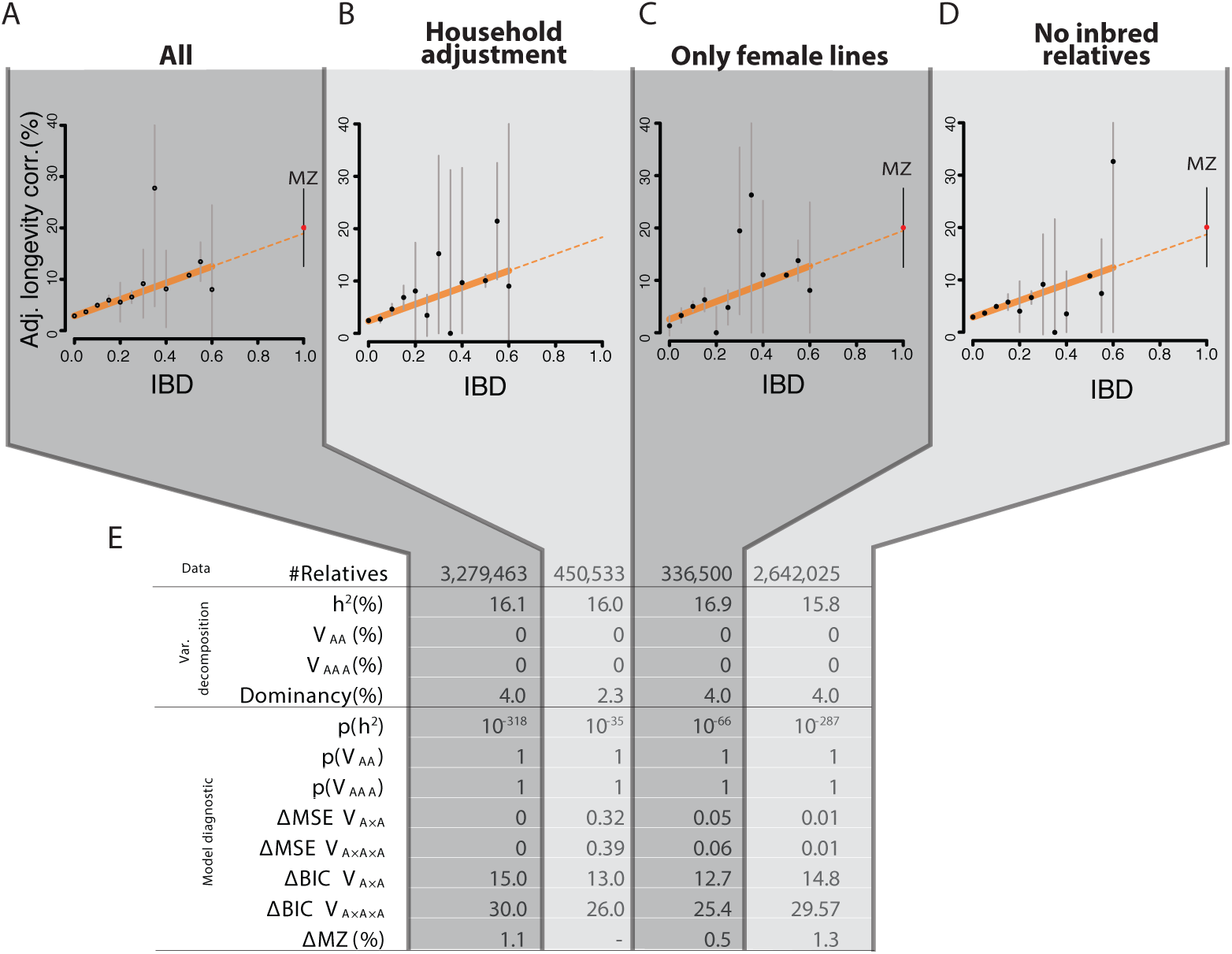
The results of variance partitioning of longevity with various adjustment of potential confounders. The figurees show the longevity correlation (after dominancy adjustment) as a function of IBD. In all cases, the epistatic terms converged to zero. The table displays the number of pairs of relatives in each condition and the maximum likelihood estimators for each component of variance. p-values denote the results of a nested ANOVA for the additive component, squared-, and cubic- epistasis. ∆MSE denotes the average residual error per sample between each epistatic model and the additive model. ∆BIC denotes BIC difference between each epistatic model and the additive model. ∆MZ shows difference between the observed correlation of longevity in the Danish MZ twins and the extrapolation of the additive model.

**Figure S13:**
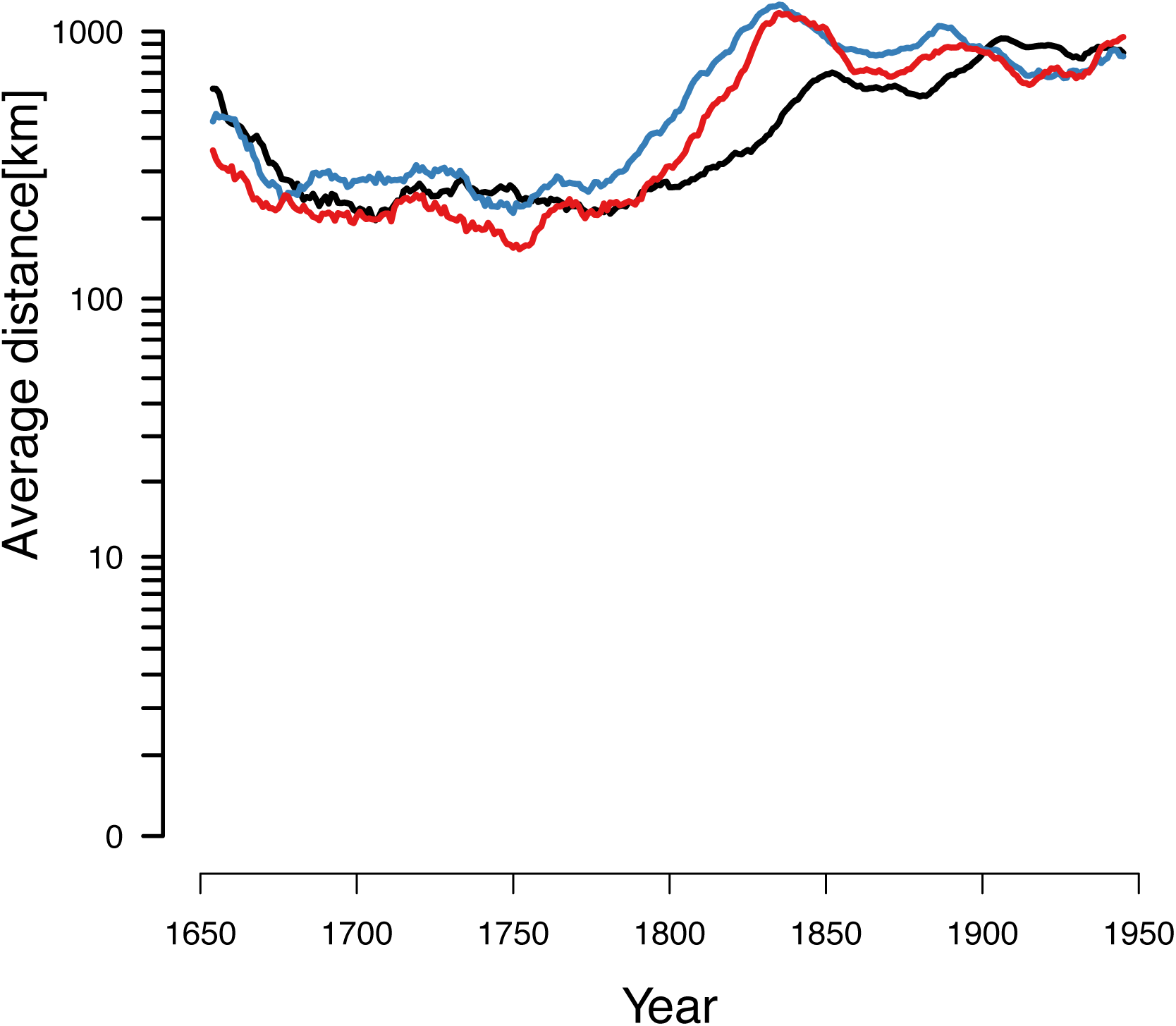
Average migration distance of individuals as a function of time. Red: mother-offspring, blue: father-offspring, black: marital radius. Dots represent the data before smoothing.

**Figure S14:**
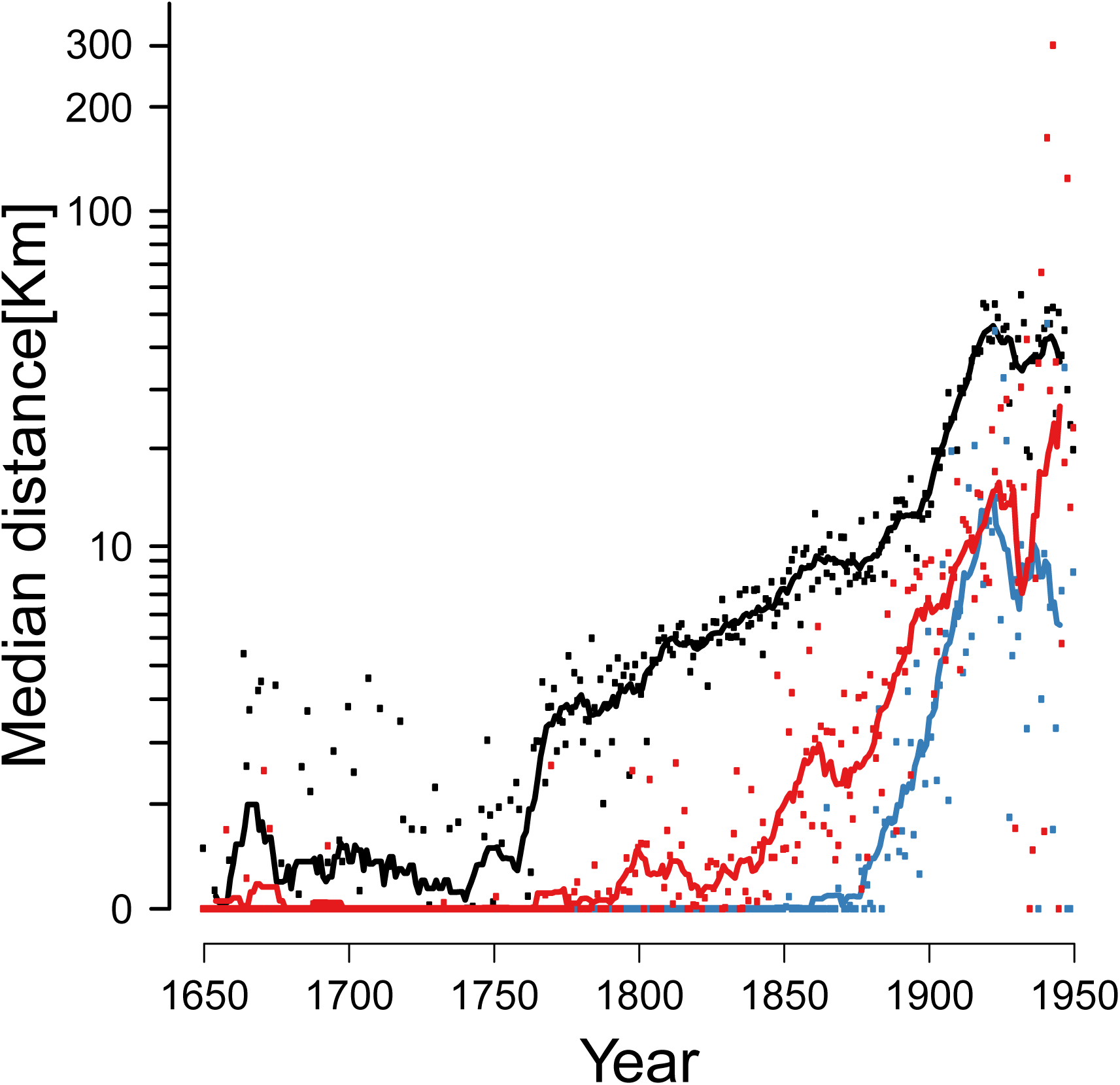
Median migration distance in European-only born individuals as a function of time. Red: mother-offspring, blue: father-offspring, black: marital radius. Dots represent the data before smoothing.

**Figure S15:**
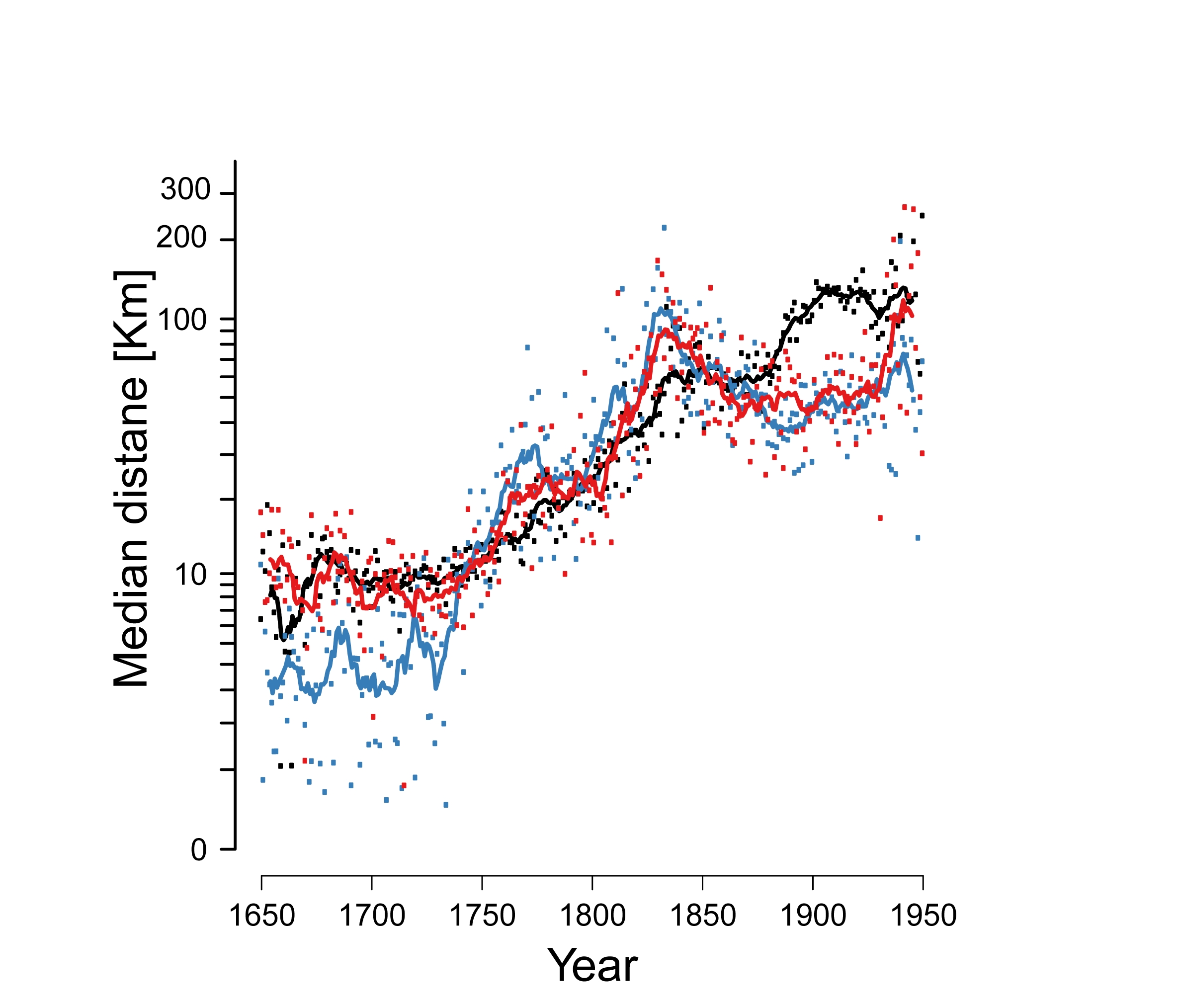
Median migration distance in North American-only born individuals as a function of time. Red: mother-offspring, blue: father-offspring, black: marital radius. Dots represent the data before smoothing.

**Figure S16:**
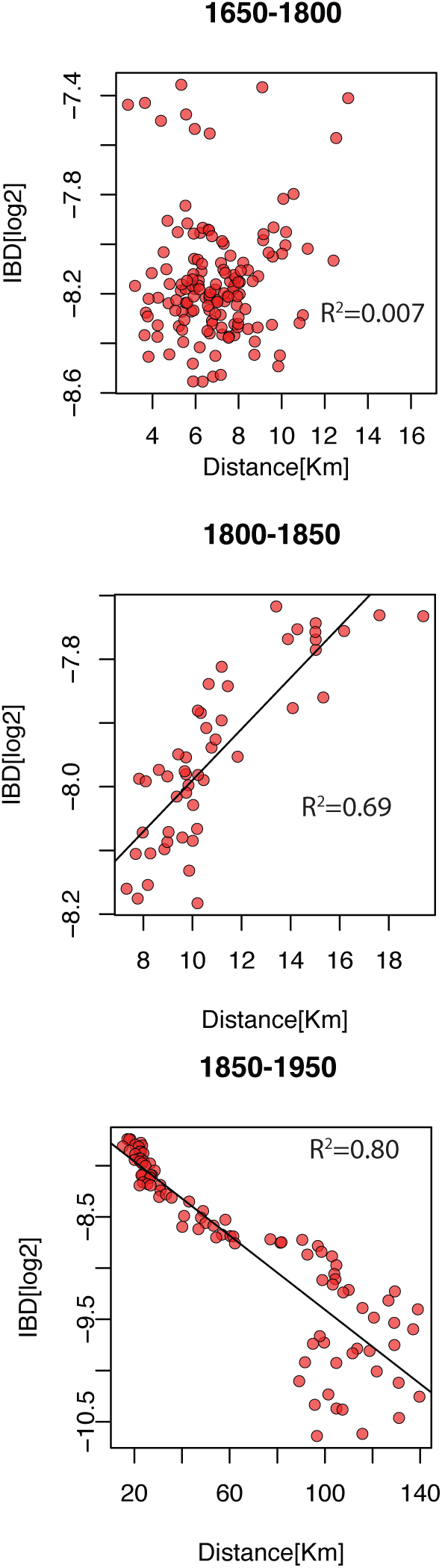
The expected kinship between couples as a function of the median martial distance stratified by time (A) Couples that were born between 1650-1800 (B) Couples that were born between 1800-1850 (C) Couples that were born between 1850-1950.

**Figure S17:**
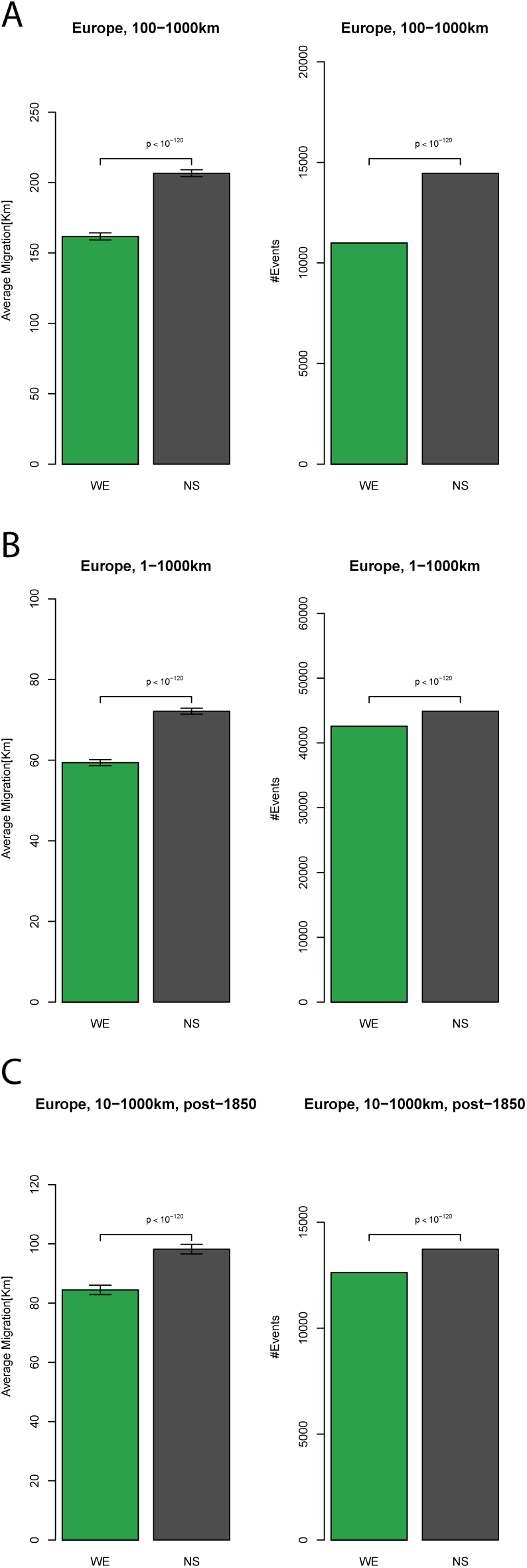
Migration direction in Europe under different conditions. Top: migration events with a distance of at least 100Km. Middle: migration events with a distance of at least 1kM. Bottom: migration events over 10Km but after 1850. Left: Difference in mean distance along the West-East axis versus North-South axis. Right: The number of events with tendency to West-East orientation versus North-South orientation.

**Figure S18:**
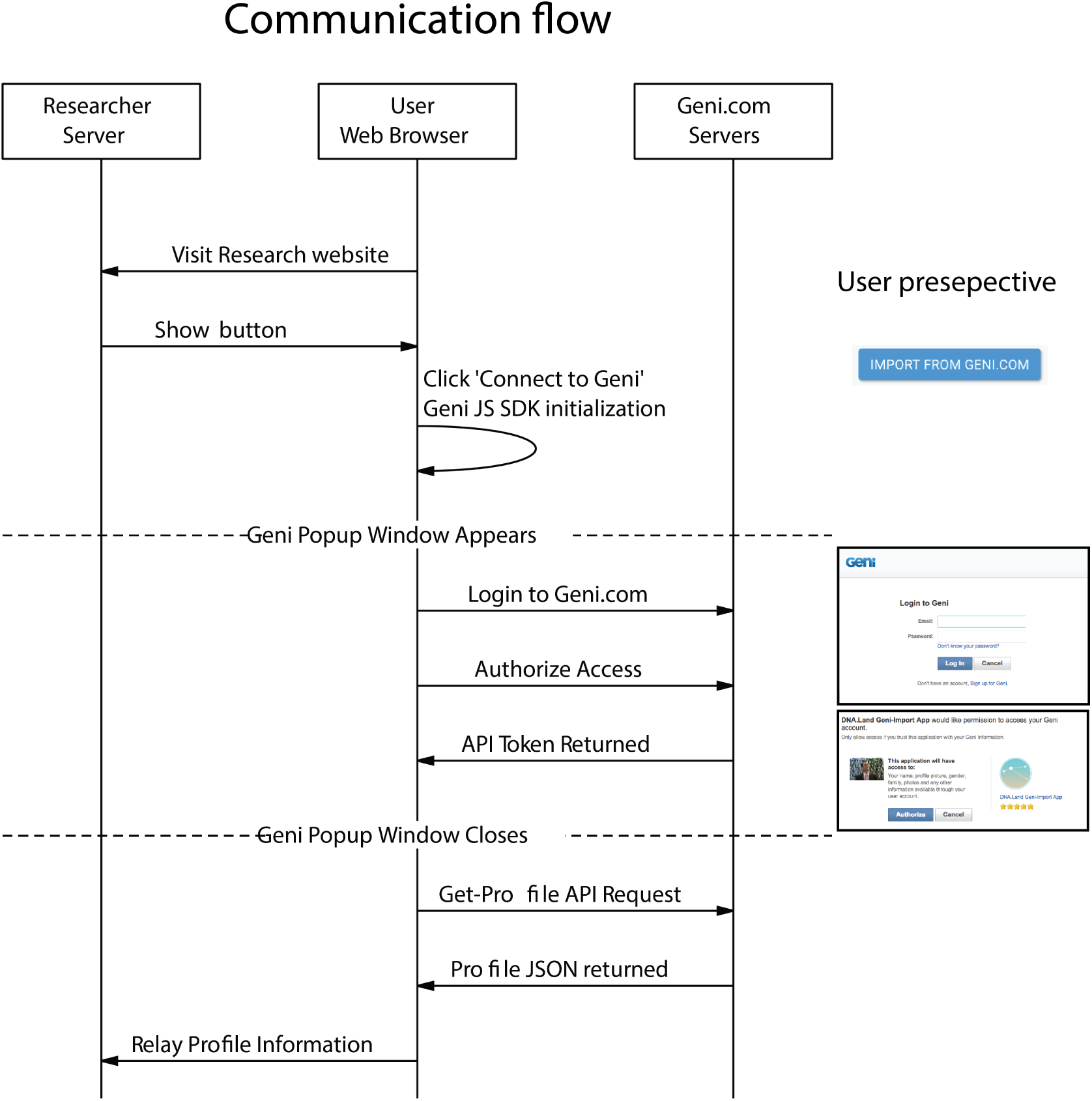
Communication flow to obtain the profile-id from users. Please see a live demo on https://teamerlich.org/geni-integration-example/

## Supplementary Tables

**Table S1:** Heritability estimates of longevity from previous studies. † For full citation, see main text.

| Paper<sup>†</sup> | Population | Relative Types | Sample Size | Years of Birth | $h^2$ estimate |
| --- | --- | --- | --- | --- | --- |
| Herskind et al. 1996 | Danish | Same sex twin pairs | 5744 | 1870-1900 | Males: 26%<br>Females: 23 |
| Mitchell et al. 2001 | Amish | Parent-offspring, siblings | 1655 | Before 1890 | 25% (20-30%) |
| Kerber et al. 2001 | Mormon | Various types up to 1st cousins | 78994 | 1870-1907<br>lived to at least 65 | 15% (12-18%) |
| Mayer 1991 | New England | 6 large families | 13656 | 1650-1874 | Parent-offspring: 10-33%<br>Sibship-Parent: 16-22%<br>Siblings: 33-41% |
| Phillipe 1978 | French Canadian | Parent-offspring | 265 | Parents married 1820-1899 | Parent-offspring: 10.1%<br>Like-sex sibling: 10.1%<br>Unlike-sex siblings: 13.9%<br>(Spouses: 12.1%) |
| Ljungquist et al. 1998 | Swedish | Same sex twin pairs | 1200 | 1886-1925 | All twin pairs: 35%<br>Male Twin pairs reared apart: 0%<br>Female Twin pairs reared apart: 15% |

**Table S2:** The number and basic properties of the pairs of relatives in FamiLinx. These are the pairs with exact date of birth, exact date of death, and exact birth location. More pairs are available but with incomplete data. Relationship: C. denotes cousinship and “R” denotes once removed. Av, G, HS, P, and S denotes avuncular, grandparent-grandchild, half-sibs, parent-offspring, and full sib relationships. 2G and 3G denote great and great-great granparent relationships. Adjusted IBD: the averaged IBD after taking into account consanguineous relationships. Distance: the median birth distance in Km. ∆Gen: the average number of years between two individuals.

| Degree of relationship | Relationship | Expected IBD | Adjusted IBD | #Pairs | Distance | ΔGen |
| --- | --- | --- | --- | --- | --- | --- |
| 1 | P | 0.5 | 0.50112 | 637,653 | 9 | 33 |
| 1 | S | 0.5 | 0.50114 | 879,694 | 0 | 7 |
| 2 | Av | 0.25 | 0.23849 | 1,500,143 | 9 | 31 |
| 2 | G | 0.25 | 0.25216 | 566,747 | 35 | 65 |
| 2 | HS | 0.25 | 0.25438 | 94,062 | 0 | 14 |
| 3 | 1C | 0.125 | 0.12281 | 1,269,077 | 9 | 11 |
| 3 | 2G | 0.125 | 0.12798 | 479,309 | 74 | 96 |
| 4 | 1RC | 0.0625 | 0.063376 | 2,573,284 | 22 | 31 |
| 4 | 3G | 0.0625 | 0.065567 | 400,327 | 123 | 128 |
| 5 | 2C | 0.03125 | 0.032895 | 2,079,209 | 30 | 14 |
| 5 | 4G | 0.03125 | 0.033906 | 322,311 | 225 | 158 |
| 6 | 2RC | 0.015625 | 0.017622 | 4,424,847 | 46 | 31 |
| 6 | 5G | 0.015625 | 0.017573 | 235,245 | 435 | 188 |
| 7 | 3C | 0.0078125 | 0.0091457 | 3,182,460 | 55 | 16 |
| 8 | 3RC | 0.0039062 | 0.005657 | 6,759,744 | 72 | 32 |
| 9 | 4C | 0.0019531 | 0.0028056 | 4,168,540 | 75 | 19 |

**Table S3:** The number of pairs and correlation of longevity for each class of relatives. The correlation was adjusted to remove dominant effects and used four cousins as the baseline.

| Relationship | Pairs of analysis | Adj. longevity correlation |
| --- | --- | --- |
| S | 253076 | 0.09231052 |
| HS | 24243 | 0.04675896 |
| 1C | 357139 | 0.02361952 |
| 2C | 584856 | 0.01122288 |
| 3C | 891462 | 0.00684701 |
| 4C | 1168687 | 0 |

